# Microglia and myeloid cell populations of the developing mouse retina

**DOI:** 10.1101/2025.04.07.646926

**Authors:** Sage Martineau, Juan C. Valdez-Lopez, Samantha Zarnick, Jeremy N. Kay

## Abstract

Microglia make important contributions to central nervous system (CNS) development, but the breadth of their distinct developmental functions remain poorly understood. The mouse retina has been a key model system for understanding fundamental mechanisms controlling assembly of the CNS. To gain insight into where and how microglia might influence retinal development, here we identified molecularly unique myeloid cell populations that are selectively present during development, and characterized their anatomical locations. Development-specific transcriptional states were identified using single-cell (sc) and single-nucleus RNA-sequencing (RNA-seq) across multiple timepoints. Transcriptional states were validated in vivo by histological staining for key RNA and/or protein markers. Several of these development-specific myeloid populations have been described before in brain scRNA-seq atlases but not validated in vivo, while others are unique to our retinal dataset. We identify two closely-related microglial populations, labeled by the *Spp1* and *Hmox1* genes, that are distinguished mainly by activity of the NRF2 transcription factor. Both types are present selectively within the developing retinal nerve fiber layer where they engulf neurons and astrocytes undergoing developmental cell death. *Hmox1^+^* microglia were also localized selectively at the wavefront of developing vasculature during retinal angiogenesis, suggesting that developmental events associated with angiogenesis modulate NRF2 activity and thereby induce microglia to switch between the *Spp1^+^* and *Hmox1^+^* states. Overall, our results identify transcriptional profiles that define specific populations of retinal microglia, opening the way to future investigations of how these programs support microglial functions during development.

## INTRODUCTION

As the tissue-resident macrophages and primary immune cells of the central nervous system (CNS), microglia play an essential role in maintaining neurological homeostasis in response to injury or disease. In recent years, there has been an increasing appreciation that microglia also have critical functions in the uninjured nervous system – especially during CNS development. Myeloid precursors colonize the CNS during early embryogenesis and are therefore positioned to influence a diverse range of developmental functions, including brain histogenesis, apoptotic cell clearance, synaptic pruning, and axon myelination (Anderson et al., 2019; Fourgeaud et al., 2016; Giera et al., 2018; Hammond et al., 2018; Lawrence et al., 2024; Wlodarczyk et al., 2017). However, in many of these cases, there remains a lack of clarity as to the specific cellular and transcriptional mechanisms by which microglia facilitate these key developmental events. Understanding such mechanisms will be important to gain a deeper insight into the role of microglia in the normal, uninjured CNS, and how microglia contribute to assembly of the nervous system.

Single-cell RNA-sequencing (scRNA-seq) has been a revolutionary tool for characterizing the transcriptional and functional diversity of brain myeloid cells, including microglia and macrophages. Numerous studies have revealed that microglia adopt well-defined transcriptional states, driven by functional needs that arise within their local environment. In adult homeostatic CNS, microglia are a homogenous population: virtually all cells express the same transcriptional profile (Chen and Colonna, 2021). By contrast, in the developing brain, scRNA-seq studies have revealed that microglia adopt numerous distinct transcriptional states, which likely reflect the variety of functions carried out by microglia to facilitate neural development (Hagemeyer et al., 2017; Hammond et al., 2019; Li et al., 2019). Some of these transient transcriptional states have been linked to a well-characterized developmental function: the phagocytic clearance of apoptotic cells generated through naturally-occurring developmental cell death (Anderson et al., 2022; Anderson et al., 2019; Hammond et al., 2019; Li et al., 2019). Microglial state(s) defined by high expression of genes such as *Itgax, Csf1, Clec7a,* and *Spp1* were shown to mark microglia that possess a classic phagocytic morphology – i.e. amoeboid or rounded rather than the highly branched or ramified morphology typical of homeostatic microglia. Moreover, scRNA-seq data revealed that these phagocytic developmental states are highly similar to the phagocytic “disease-associated microglia” state induced by a variety of pathological insults in adult CNS (Anderson et al., 2022; Anderson et al., 2019; Benmamar-Badel et al., 2020; Ghena et al., 2025; Krasemann et al., 2017; Li et al., 2019). While these studies have begun to link developmental phagocytic activity to particular microglial populations, there are many development-specific transcriptional states whose functions remain uncharacterized. Furthermore, most studies so far have focused on specific brain regions such as the cerebral cortex, corpus callosum, and visual thalamus; developmental studies of microglia within other CNS regions are needed to learn whether there are region-specific microglial populations that carry out specialized developmental functions.

Here we provide a comprehensive profile of microglia and macrophage diversity within the developing mouse retina – a long-standing model system for studies of neural development. To understand development-specific functions of microglia, one particular advantage of the retina relative to the brain is its highly laminated structure, with neuronal cell bodies, synapses, axons, and vasculature all occupying distinct retinal layers (Fig. 1A). Thus, association of microglia with particular cell types or neuronal subcellular compartments can be inferred from their laminar location. In uninjured adult retina, microglia selectively occupy the inner and outer plexiform layers (IPL and OPL), indicating that synaptic structures provide a niche for homeostatic microglia. By contrast, the laminar distribution of microglia is more diverse during development (Puñal et al., 2019), suggesting that transient developmental events necessitate recruitment of microglia to other retinal layers. One prominent example is formation of the primary vascular plexus within the retinal nerve fiber layer (RNFL; Fig. 1A). Starting at postnatal day (P) 0, an orderly wave of angiogenesis spreads outward from the optic nerve head through the RNFL, following a two-dimensional template comprised of astrocyte arbors. Microglia populate the RNFL during this process, where they influence angiogenesis in multiple ways – including by promoting developmental death/remodeling of the RNFL astrocyte population (Kubota et al., 2009; Puñal et al., 2019; Tammela et al., 2011). How microglia accomplish these pro-angiogenic functions is not well understood, but important insights into this problem could emerge from determining the identity and molecular characteristics of these transient RNFL microglia.

**Figure 1:**
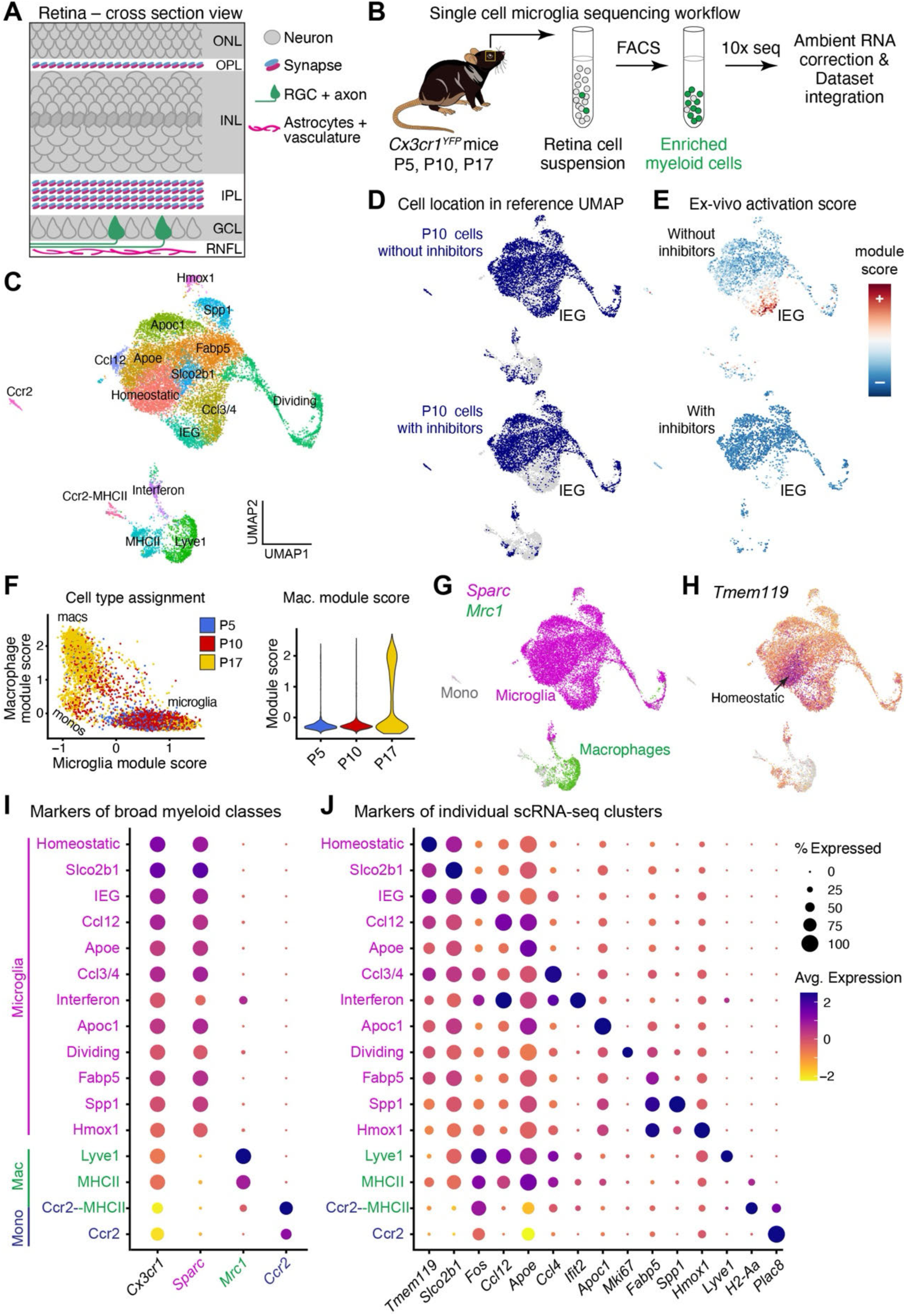
Profiling myeloid cell transcriptional diversity across retinal development. **A.** Illustration showing cellular contents of retinal layers. ONL/INL, outer/inner nuclear layer; other abbreviations, see main text. **B.** Workflow for isolation and single-cell sequencing of retinal myeloid cells at P5, P10, and P17 (also see Methods). **C.** UMAP plot generated from the integrated P5-P10-P17 dataset. 17,338 myeloid cells are assigned to 16 clusters via unsupervised clustering. **D,E**. Effects of transcriptional + translational inhibitor cocktail on scRNA-seq gene expression. (D) UMAP locations (blue) of cells from P10 datasets generated without (top) or with (bottom) inhibitors. Inclusion of inhibitors selectively depleted cells from the IEG cluster. (E) Inhibitors markedly reduced expression of a transcriptional profile associated with ex-vivo gene activation. **F.** Classification of cells as microglia, macrophages (macs), or monocytes (monos) using module score scatterplot. Colors indicate age metadata for each cell. While some macrophages were detected at P10 (left), they only became an appreciable fraction of retinal myeloid cells at P17 (right). **G.** Each myeloid cell class occupies distinct regions of the UMAP plot, as shown by expression of microglia marker *Sparc* (magenta) and macrophage marker *Mrc1* (green). Monocytes (mono) express neither marker. **H.** High expression of *Tmem119* distinguishes homeostatic microglia cluster (arrow). **I.** Dot plot showing expression of markers for broad myeloid cell classes across clusters. Magenta, microglia; green, macrophages; blue, monocytes. **J.** Dot plot showing expression of a selected top marker of each cluster.

To comprehensively survey microglial diversity within developing mouse retina, we performed single-cell sequencing of over 17,000 myeloid cells isolated at three timepoints spanning postnatal retinal development. This dataset identified numerous microglia and macrophage populations showing distinct transcriptional profiles. We then stained ocular tissue with RNA and/or protein markers for many of these transcriptional states, thereby identifying microglial and macrophage populations that are selectively present at specific times and/or places within the developing eye. We identify a microglial population that is selectively associated with RNFL development, where its functions include removal of astrocytes as well as apoptotic neurons from the adjacent ganglion cell layer (GCL). Further, we show that this RNFL population comprises two inter-related transcriptional states that, based on scRNA-seq studies in the developing brain, were previously thought to be separate (Hammond et al., 2019). Altogether, our results provide a foundation for future studies into development-specific myeloid cell functions, both within the visual system and across the CNS.

## RESULTS

### Generation of a myeloid scRNAseq dataset from developing mouse retina

To characterize the diversity of microglia and other myeloid cells in developing mouse retina, we performed scRNAseq at three timepoints – postnatal day (P)5, P10, and P17 (Fig. 1B). The first two timepoints were chosen to capture different stages of postnatal retinal development, whereas the third timepoint represents an age at which development is largely complete. Retinas were isolated from *Cx3cr1^YFP^* animals (Parkhurst et al., 2013) and YFP^+^ myeloid cells were purified by flow cytometry. These samples were then processed for 10x Genomics scRNAseq, corrected for ambient RNA contamination (Young and Behjati, 2020), and analyzed using standard single-cell bioinformatics approaches (Stuart et al., 2019). After filtering cells that did not pass sequencing quality control metrics or that expressed neuronal markers (see Methods), we obtained a dataset containing 17,338 total myeloid cells (3,920 from P5, 8,995 from

P10, and 4,423 from P17). Data from independent replicates and timepoints were integrated, clustered, and a UMAP projection was generated to identify putative cellular populations (Fig. 1B,C). This workflow initially identified 18 clusters. Three of these corresponded to cells in various stages of the cell cycle; these were combined into a single dividing-cell cluster, leaving us with 16 total clusters (Fig. 1C). The vast majority of cells expressed markers of either macrophages, monocytes, or microglia, confirming that contaminants had been successfully eliminated and that the clusters reflect transcriptional diversity of myeloid cells (Fig. 1F,G; Supplemental Fig. S1).

We next sought to validate that the scRNA-seq clusters reflect bona fide gene expression states of myeloid cells in vivo. To this end, we conducted additional sequencing experiments designed to identify and minimize ex vivo gene expression artifacts. First, to eliminate expression artifacts that arise during tissue dissociation, we performed single-nucleus (sn) RNA-seq on nuclei isolated from P5 and P10 flash-frozen retinal tissue. Nuclear envelopes of *Cx3cr1*^+^ myeloid cells were genetically labeled (Mo et al., 2015) and enriched by flow cytometry (Supplemental Fig. S2A). Subsequently, P5 and P10 snRNA-seq datasets were generated using a bioinformatics workflow similar to the scRNA-seq studies, and the single-nucleus data were projected onto the integrated scRNA-seq UMAP to facilitate comparisons between the two datasets (Supplemental Fig. S2B). This analysis revealed that snRNA-seq successfully diminished ex vivo artifacts: the scRNA-seq cluster expressing high levels of immediate-early genes (IEGs) was absent from single-nucleus data, consistent with prior work showing that this cluster arises due to ex vivo IEG activation during tissue dissociation (Marsh et al., 2022) (Supplemental Fig. S2). However, we also noted – consistent with prior reports (Thrupp et al., 2020) – that certain other clusters were also missing from the nucleus dataset, owing to their key marker genes being poorly detected by snRNA-seq (Supplemental Fig. S2B,C). This observation suggests that the snRNA-seq protocol might have a bias against detecting certain microglial states.

We therefore sought an alternative approach for reducing ex vivo expression artifacts. To this end, we combined scRNA-seq with inhibition of ex vivo gene expression. A single-cell sample of FACS-sorted *Cx3cr1^YFP^*^+^ myeloid cells was prepared at P10 using a cocktail of transcriptional and translational inhibitors (Marsh et al., 2022), and the resulting scRNA-seq dataset was compared to the P10 dataset generated without this inhibitor cocktail. We found that inclusion of inhibitors dramatically reduced the number of cells assigned to the IEG cluster; by contrast, all other clusters contained cells from the inhibitor-treated sample (Fig. 1D). Furthermore, none of the microglia from the inhibitor sample showed high expression of the artifactual ex-vivo gene activation profile that has been described in previous studies (Fig. 1E) (Marsh et al., 2022). These observations suggest that the IEG cluster is the only one in our scRNA-seq dataset representing an obvious ex vivo activation state, whereas the other transcriptional profiles captured by this dataset are likely to exist in vivo.

### Overview of retinal myeloid cell populations identified by scRNA-seq

To document the cell types and cell states corresponding to the non-IEG scRNA-seq clusters, we first examined expression of known marker genes for the broad myeloid cell classes. This analysis revealed two major subdivisions of the UMAP plot corresponding to *Sparc1^+^*microglia and *Mrc1^+^* macrophages, along with a small *Ccr2*^+^ monocyte population (Fig. 1G,I). Subsequently, as a more robust way of assigning cell type identity, we used a module score approach (Tirosh et al., 2016) which enables cellular phenotyping using a list of cell type marker genes rather than a single gene.

Using microglia and macrophage module scores (Supplemental Table S2), most cells could be unambiguously assigned to one of the three broad myeloid cell classes (Fig. 1F). Only two clusters – the Dividing cluster and a cluster defined by high expression of interferon-related genes – contained mixtures of both microglia and macrophages; other clusters were largely comprised of just one of these cell types (Fig. 1I). Macrophages and monocytes were not detected in appreciable numbers until P17, suggesting that they become established relatively late in retinal development (Fig. 1F).

We next assessed whether the clusters correspond to known myeloid cell types and/or transcriptional states. To this end, differential gene expression analysis was used to identify the top marker genes for each cluster (Fig. 1J; Supplemental Table S1).

Clusters with a clear correspondence to previously described cell types/states were given the same name as in prior studies; otherwise, clusters were named based on their top marker gene (cluster names are capitalized and non-italic; gene names are italicized per mouse gene naming convention). Among microglia, one cluster clearly corresponded to the homeostatic “resting” microglia state that is observed in adult retina and brain (Hammond et al., 2019; O’Koren et al., 2019), based on high expression of known homeostatic markers including *Tmem119* (Fig. 1H,I) (Chen and Colonna, 2021). The remaining microglial clusters appeared to be in various alternative transcriptional states, based on 1) lower expression of homeostatic markers such as *Tmem119*; and 2) strong expression of unique marker genes that were not highly expressed by the putative homeostatic microglia (Fig. 1H,J).

The non-microglial populations comprised four separate clusters: one containing monocytes, two containing macrophages, and one containing a mixture of monocyte-derived cells (Fig. 1I). Based on their top markers, we matched each of these populations to myeloid cells described in previous scRNA-seq studies. The Ccr2 cluster expresses markers of *Ear2^+^* monocytes – a Ly6C^low^ monocyte cell type that has been detected in adult mouse eyecup (Supplemental Fig. S1) (Benhar et al., 2023). The two macrophage clusters, Lyve1 and MHCII, express multiple markers of border-associated macrophages (BAMs) that have been characterized in brain (Fig. 1I,J; Fig. 2A; Supplemental Fig. S1) (Brioschi et al., 2023; Utz et al., 2020; Van Hove et al., 2019).

**Figure 2:**
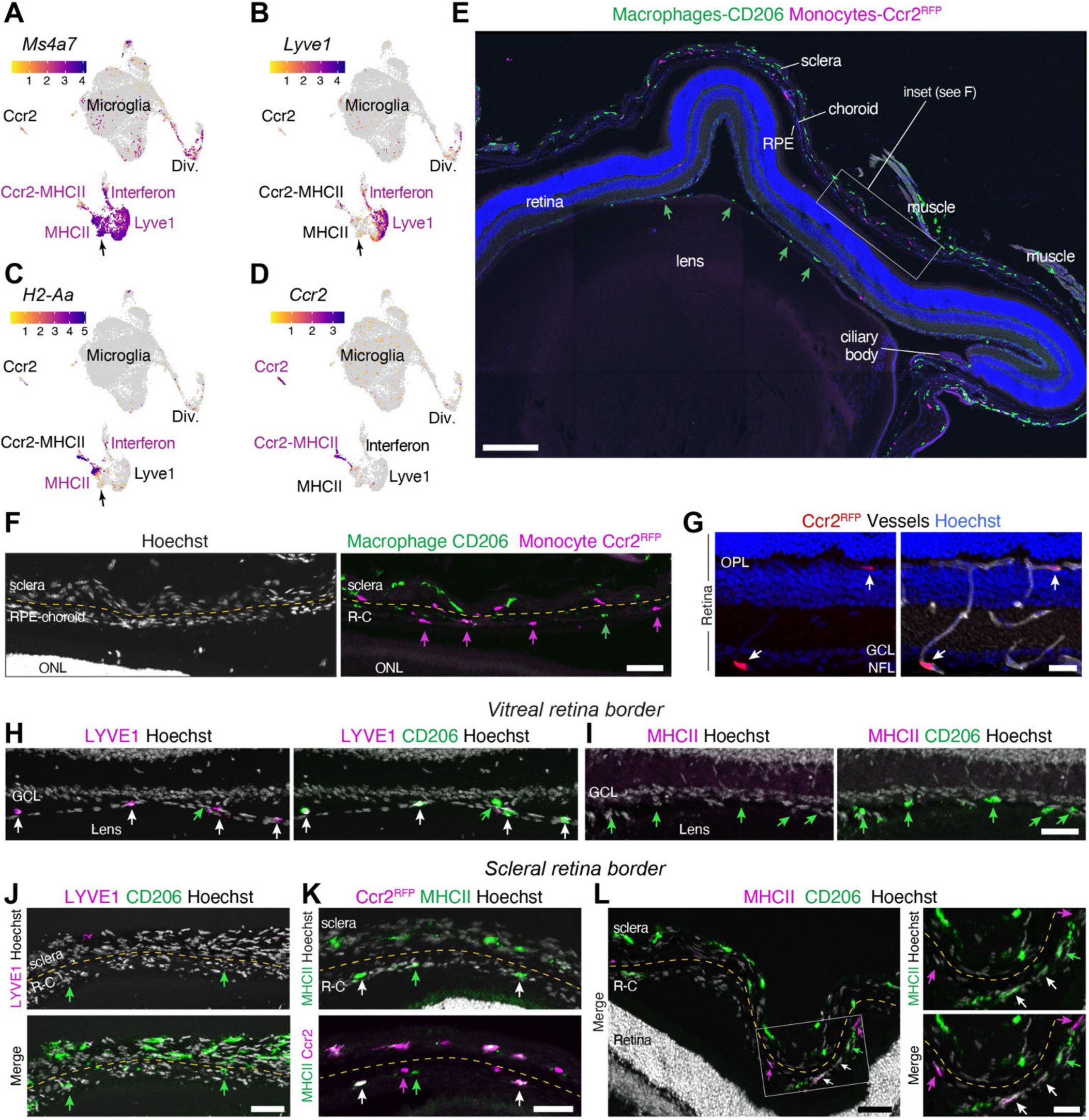
Characterization and validation of macrophage and monocyte clusters. **A-D.** Expression of markers for defined macrophage and monocyte populations that were assessed histologically. *Ms4a7*, pan-BAM marker, similar to *Mrc1* (Fig. 1G,I). *Lyve1* and *H2-Aa*; BAM subtypes. *Ccr2*, monocytes. Clusters with large fractions of cells expressing each marker are denoted in magenta text. The Interferon cluster contained both BAM subtypes. A subdivision of the MHCII cluster (arrow) corresponds to *Ms4a7^+^* BAMs that do not express *H2-Aa* or *Lyve1* (see also Supplemental Fig. S1). **E**. P17 whole-eye sections immunostained for monocytes (Ccr2-RFP) and macrophages (CD206, encoded by *Mrc1* gene). Both cell types are detected in sclera, RPE, choroid, and ciliary body (cb). CD206 also labels macrophages adhering to lens and inner retinal surface (green arrows), likely hyalocytes. Monocytes are absent from vitreous/lens region. **F**. Higher-magnification view of boxed region in E showing mutually exclusive expression of Ccr2-RFP and CD206 at outer retina border (right panel). Green arrows, CD206^+^ cells; magenta arrows, Ccr2-RFP^+^ cells. Hoechst nuclear stain (left panel) was used to identify RPE, choroid and sclera layers. In some regions, distinguishing RPE from choroid was difficult due to tissue compression during sectioning, so they were labeled together. A clear gap was always present between the choroid and sclera (dashed line). **G**. Retinal Ccr2-RFP monocytes (arrows) are located inside vasculature (IB4 lectin, white), not within retinal parenchyma (Hoechst, blue). **H,I**. CD206^+^ BAMs at vitreal retina border are immunoreactive for LYVE-1 (H, magenta) and do not express MHCII (I, magenta). White arrows, double-labeled cells. Green arrows, CD206^+^ single-labeled cells without the magenta marker. **J**. LYVE-1 expression is absent from outer retina border tissues. Green arrowheads, CD206^+^ cells within the RPE-choroid (R-C) layers that lack LYVE-1. **K**. Cells co-expressing Ccr2-RFP and MHCII (white arrows) are detected in RPE, choroid, and sclera. These likely correspond to the Ccr2-MHCII cluster. Green arrows, MHCII^+^ cells without Ccr2. **L**. CD206^+^ BAMs within the RPE-choroid layers comprise two populations: MHCII^+^ CD206^+^ double-positive cells (white arrows) and MHCII^−^ CD206^+^ cells (green arrows). As these cells are also LYVE-1^−^ (J), they likely correspond to two populations of cells within the MHCII cluster (C, arrow). MHCII^+^ cells that lack CD206 (magenta arrows) are most likely the Ccr2^+^ monocyte-derived population (see F,K). Panels at right show high-magnification view of boxed region. All images are representative of multiple sections imaged from n = 3 separate mice. Scale bars, 200 µm (E); 50 µm (F, H-K, L left panel); 25 µm (G, L right panels).

Further supporting their classification as the ocular counterparts to BAMs, the two clusters individually express markers of well-characterized cranial BAM subpopulations – one marked by *Lyve1* and the other marked by major histocompatibility class II (MHCII) components such as *H2-Aa* (Fig. 1J; Fig. 2B,C; Supplemental Fig. S1). Finally, the Ccr2-MHCII cluster expresses markers of monocyte-derived cells of multiple types (Supplemental Fig. S1). Many cells in this cluster co-express markers of the MHCII^+^ BAMs alongside markers of monocytes, suggesting that this population might be differentiating from monocytes into macrophages (Fig. 1I,J; Fig. 2D; Supplemental Fig. S1). This possibility is consistent with previous findings that brain MHCII^+^ BAMs are mainly monocyte derived (Brioschi et al., 2023). Another subset of cells within the Ccr2-MHCII cluster likely represents monocytes differentiating into dendritic cells, based on the co-expression of *H2-Aa, Ccr2,* and *Cd209a* (Supplemental Fig. S1) (Benhar et al., 2023).

We conclude from this initial analysis that there is substantial diversity among myeloid cells in developing retina, and that our dataset provides a resource for defining the transcriptional profiles of these diverse cell states.

### Border associated macrophage subtypes localize to distinct ocular tissues

Having completed construction of a retinal myeloid scRNA-seq dataset, we next sought to experimentally validate that these correspond to valid cell types in vivo. We began by characterizing the macrophage and monocyte populations (Fig. 2A-D). Even though our dataset was produced using retinal tissue that had been dissected free of the eyecup and anterior ocular tissues, we hypothesized that the non-microglial populations most likely derive from tissues bordering the neural retina rather than the retina itself. This idea was motivated by the similarity of the Lyve1 and MHCII macrophages to cranial BAMs, which do not populate the brain parenchyma but instead localize to brain borders (Munro et al., 2022). Ocular border tissues that could plausibly be sources of monocytes or macrophages, based on how tissue was collected, include the vitreous, choroid, and retinal pigment epithelium (RPE), as well as the perivascular space. Cells from these regions could have adhered to the retina during tissue isolation. By contrast, cells from the sclera or anterior eye structures, such as the cornea and limbus, are unlikely to be represented in our dataset.

To determine the tissue localization of *Ccr2*^+^ monocytes and *Mrc1^+^* macrophages in vivo, we performed immunohistochemistry on sections prepared from whole P17 mouse eyes. *Ccr2^+^*monocytes were labeled using a well-characterized *Ccr2^RFP^* knock-in mouse line, in which red fluorescent protein (RFP) is expressed from the *Ccr2* locus (Saederup et al., 2010). BAMs were labeled by antibodies to CD206, the protein encoded by *Mrc1*. This analysis revealed selective localization of both cell types to regions bordering the retina, whereas the retina itself was devoid of RFP^+^ or CD206^+^ cells (Fig. 2E). This included an absence of perivascular macrophages adjacent to intraretinal vessels, suggesting that such cells are either not yet present or do not yet express CD206 by P17. While occasional Ccr2-RFP^+^ monocytes were noted within the retina, these were circulating through intraretinal vessels and were not residents of the CNS (Fig. 2G).

In contrast to the retina itself, regions bordering the outer and inner retina exhibited prominent labeling for Ccr2-RFP*^+^* monocytes and/or CD206^+^ macrophages. At the outer retinal border, prominent and mutually exclusive Ccr2-RFP and CD206 labeling was observed within the choroid and RPE (Fig. 2E,F). A similar pattern was also observed in the sclera, although such cells were unlikely to be present in the scRNA-seq tissue preparation. The lack of overlap between RFP and CD206 indicates that ocular monocytes and macrophages are distinct cellular populations (Fig. 2E,F). At the inner retinal border, CD206 labeled a population of macrophages, which likely includes the hyalocytes (Rajesh et al., 2022; Rosmus et al., 2024), adhering to the lens and the vitreal surface of the retina (Fig. 2E,H,I). RFP^+^ monocytes, by contrast, were absent from the vitreal region (Fig. 2E). These findings indicate that CD206^+^ BAMs are present within tissues bordering both the vitreal and scleral side of the retina, while monocytes are restricted to scleral-side border tissues such as the RPE and choroid.

We next sought to determine the ocular location of the Lyve1 and MHCII BAM subpopulations predicted by our dataset (Fig. 2B,C). Antibody staining for LYVE-1 and the MHCII complex revealed striking tissue specificity: LYVE-1^+^ macrophages localized selectively to the vitreous and lens, whereas MHCII^+^ macrophages were found exclusively within scleral-side border tissues (Fig. 2H-L). This latter population included MHCII^+^ CD206^+^ BAMs, corresponding to the MHCII cluster in our dataset, as well as MHCII^+^ Ccr2-RFP^+^ cells corresponding to the Ccr2-MHCII cluster (Fig. 2K,L; Supplemental Fig. S1). Occasional CD206^+^ cells lacking MHCII were also observed in the RPE and choroid, suggesting that an additional BAM population, which lacks LYVE-1 and MHCII expression, might be present at the scleral retinal border (Fig. 2L). This possibility is supported by our sequencing data: Within the MHCII cluster, there was a subset of cells that only minimally expressed MHCII genes but were otherwise transcriptionally similar to the MHCII^+^ cells in the cluster (Fig. 2C, arrow; Supplemental Fig. S1; Van Hove et al., 2019). These may constitute a distinct BAM subtype that clustered with MHCII^+^ cells; or else they may be MHCII-type BAMs that are transiently expressing low levels of MHCII genes. Either way, our staining suggests that this LYVE-1^−^ MHCII^−^ CD206^+^ population exists within the RPE-choroid region. Together, these data demonstrate the existence of multiple BAM subtypes bordering the mouse retina, which express different transcriptomic phenotypes depending on whether they are located within vitreal or scleral border tissues.

### Transcriptional diversity of developing retinal microglia

We next interrogated the diversity of microglia within our dataset. To this end, we filtered out other myeloid cell types (i.e. macrophages and monocytes) and re-ran the scRNAseq bioinformatics workflow. Unsupervised clustering of the filtered dataset identified 16 microglial clusters, which were again named either by their top marker genes or based on similarity to previously described scRNAseq clusters (Fig. 3A; Supplemental Fig. S3). Most of the clusters were recognizable from the full dataset (Fig. 1B); however, four new clusters were also detected: 1) a second cluster expressing a distinct set of interferon-responsive genes (Ifi27l2a); 2) a small cluster expressing arginase (*Arg1*) as its top marker; 3) a new subdivision of the original Apoe cluster, marked by *Ccdc152*; and 4) a cluster marked by *Cdkn1a*, which encodes the p21 cyclin-dependent kinase inhibitor and is often upregulated in response to cellular stressors (Fig. 3A; Supplemental Fig. S3; Supplemental Table S3). Cells from the inhibitor+ P10 dataset populated 14 out of the 16 microglia clusters, indicating that these transcriptional states are unlikely to be artifacts of ex vivo gene activation (Supplemental Fig. S3B). The only exceptions were the IEG cluster, which corresponds to the artifactual cluster we identified in our analysis of the full dataset (Fig. 1D,E); and the Arg1 cluster, which was exclusively detected at P5 and therefore was absent from both P10 datasets regardless of inhibitor treatment (Fig. 3B).

**Figure 3.**
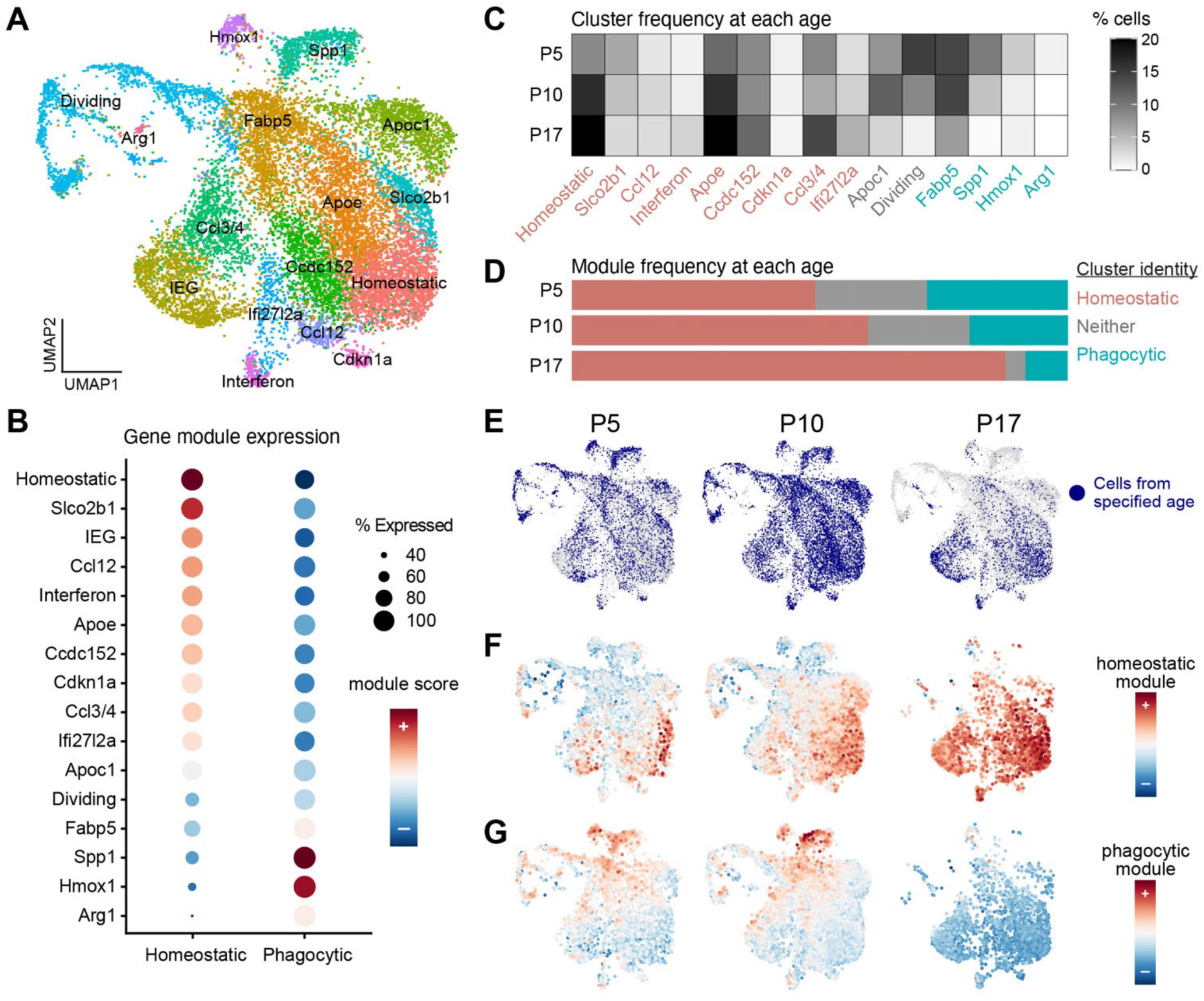
Microglia transcriptional diversity across retinal development. **A**. UMAP plot showing the integrated microglia dataset. Cluster names reflect either a top marker gene (Supplemental Fig. S2) or transcriptional similarity to previously described scRNA-seq clusters. Object was generated by removing macrophages and monocytes from each individual P5, P10, and P17 dataset, followed by reciprocal PCA integration. **B**. Dot plots showing mutually exclusive expression of homeostatic and phagocytic gene modules. Microglia clusters were classified as homeostatic or phagocytic (or neither) based on their expression of these two gene sets. **C,D.** Changes in microglia transcriptional identity across development. C: The fraction of microglia within each cluster at each timepoint. D: The fraction of microglia within clusters classified as homeostatic (red), phagocytic (aqua), or neither (gray), based on module scores (B). Phagocytic microglia diminished over time, while homeostatic microglia became more prevalent. **E**. Location of microglia from each age (blue) within the microglia UMAP plot. P5 and P10 microglia occupy all regions of the plot, while P17 microglia are predominantly located within homeostatic clusters. **F,G**. UMAP plots in which P5, P10, and P17 microglia are colored by their homeostatic (F) or phagocytic (G) module score. Homeostatic scores are higher at P17 than at earlier stages, and phagocytic module expression is largely absent by P17.

Based on their top marker genes, many of the 16 microglia clusters correspond to ones that have been observed in previous scRNA-seq studies of developing brain or retina. This group includes Homeostatic and Dividing microglia, as well as Spp1, Hmox1, Fabp5, Apoe, Ccl3/4, Ifi27l2a, and Arg1 clusters (Anderson et al., 2022; Chen and Colonna, 2021; Hammond et al., 2019; Li et al., 2019). However, there were also clusters for which we could not identify an obvious match in our review of prior studies of the homeostatic mouse CNS. These novel clusters include those marked by high expression of *Apoc1*, *Ccl12, Cdkn1a, Slco2b1*, and *Ccdc152* (Fig. 3A; Supplemental Fig. S3A).

To begin characterizing the diverse microglial states predicted by our dataset, we first asked whether any of the 16 clusters may be functionally grouped together based on shared transcriptional profiles. Prior microglia scRNA-seq studies have noted a dichotomy between two functionally distinct transcriptional states: 1) the homeostatic state, expressed by resting microglia in uninjured mature CNS; and 2) a non-homeostatic state in which microglia express a characteristic gene signature related to phagocytic functions. This phagocytic state has been observed both in developing CNS (Anderson et al., 2019; Hammond et al., 2019; Li et al., 2019) as well as in adult disease or injury contexts (Anderson and Vetter, 2019; Benmamar-Badel et al., 2020; Chen and Colonna, 2021). To determine the extent to which our 16 microglia clusters resembled these well-characterized homeostatic and phagocytic states, we examined expression of core gene modules defining each state (Supplementary Table S2; see Methods for details on module gene selection). This analysis revealed that most microglial clusters could be clearly defined as expressing either a homeostatic or phagocytic phenotype: Clusters that scored highly for one gene module scored low for the other (Fig. 3B). There were 10 clusters that upregulated the homeostatic gene module (including the Homeostatic cluster itself), while 4 clusters upregulated the phagocytic profile. Only two clusters – Apoc1 and Dividing – did not highly express either module (Fig. 3B). Thus, the diverse microglial populations within developing retina largely belong to two broad phenotypic classes defined by the homeostatic and phagocytic gene programs.

We next asked how retinal microglia diversity changed over developmental time. Whereas developing microglia (i.e. P5 and P10) populated all clusters, most P17 microglia belonged to one of the homeostatic clusters (Fig. 3C-E). Accordingly, the proportion of retinal microglia within phagocytic clusters diminished over time (Fig. 3D,E). Furthermore, regardless of their unsupervised cluster assignment, P17 microglia universally upregulated the homeostatic gene module while downregulating the phagocytic module (Fig. 3F,G). Indeed, microglia expressing the phagocytic module were largely absent by P17 (Fig. 3G). Together, these results indicate that retinal microglia are more diverse during development than in adults; and that this loss of diversity is driven by a shift from non-homeostatic to homeostatic transcriptional states.

### Characterization of microglia states specific to developing retina

From our scRNA-seq dataset we identified multiple microglial clusters that were enriched in developing retina, thereby defining transcriptional states that could subserve development-specific microglial functions (Fig. 3C,E). To better understand these developmental states, and to validate their existence in vivo, we performed additional bioinformatics analysis and stained retinal tissue with cluster-specific markers. We focused for these studies on the five clusters that showed the largest changes in population size across development. This group included all four clusters that expressed the phagocytic transcriptional module – Arg1, Fabp5, Spp1, and Hmox1 – as well as the Apoc1 cluster.

<u>Apoc1 cluster:</u> The cluster marked by expression of the lipoprotein-encoding gene *Apoc1* (Fig. 4A,B; Supplemental Fig. S3A) was transcriptionally distinct from both homeostatic and phagocytic microglia (Fig. 3B). While the cluster was distinguished by numerous quantitative differences in expression of particular genes, *Apoc1* itself was the only definitive marker gene for this cluster, showing 6-fold enrichment compared to other microglia in the dataset (Supplemental Table S3). By contrast, the second-best marker of this cluster, *Lyz2*, was only ∼1.4-fold higher (Supplemental Table S3). Transcriptionally, this cluster resembles *Apoc1*-expressing microglial clusters identified in previous single-cell studies; however, a major difference from those studies was that they only detected *Apoc1^+^*microglia in brain disease states or following microglial ablation (Anderson et al., 2022; Masuda et al., 2019). By contrast, we detected *Apoc1^+^* microglia in the uninjured developing retina. To our knowledge, these cells have not previously been detected under homeostatic conditions or during normal CNS development.

**Figure 4.**
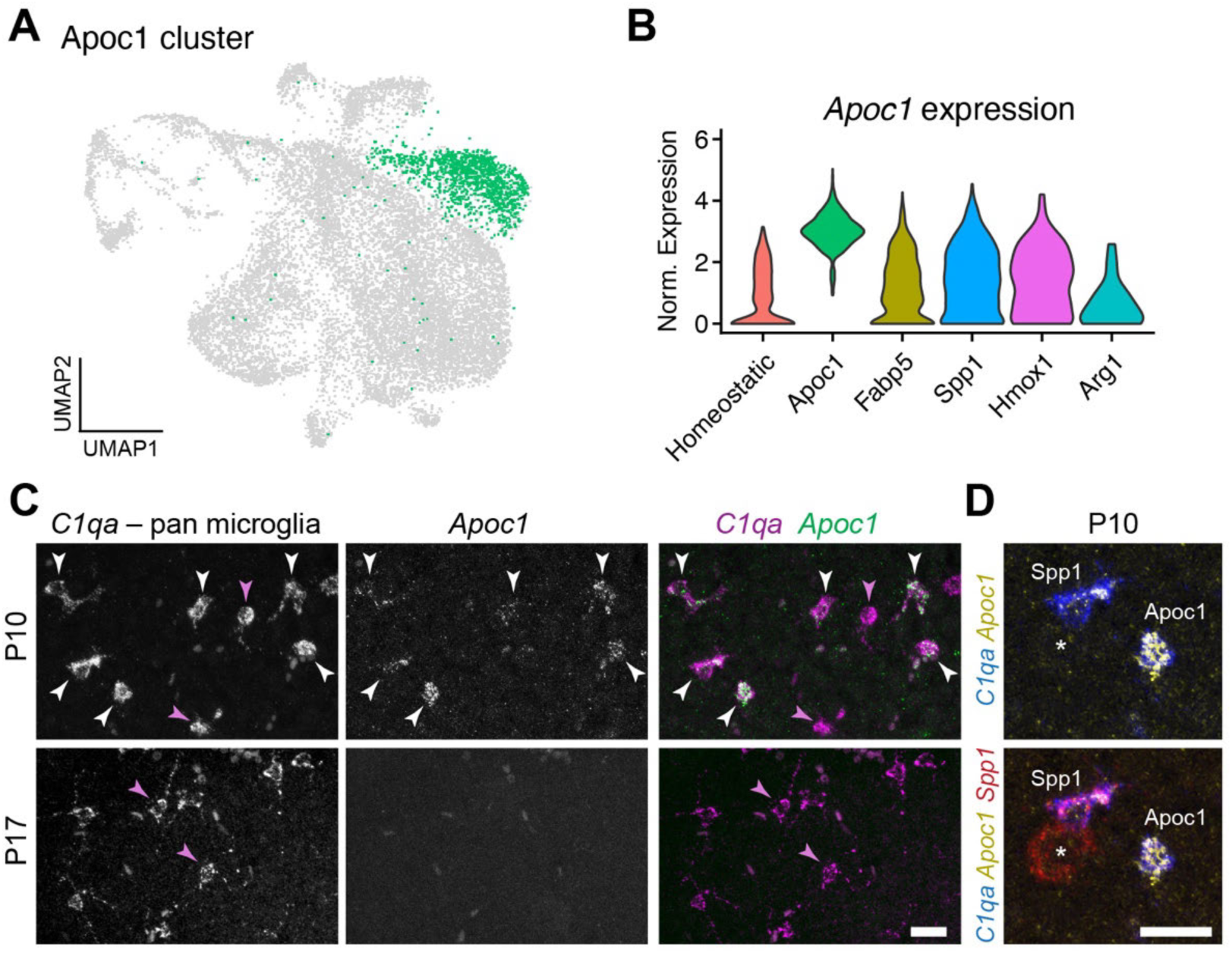
Characterization of Apoc1 and Arg1 microglial states. **A.** Apoc1 cluster highlighted on the microglia UMAP plot (see Fig. 4A). **B.** Violin plot showing *Apoc1* expression in the 5 development-specific clusters and the homeostatic cluster. Some *Apoc1*^+^ cells are present in other clusters, but *Apoc1* expression levels are far lower than for microglia assigned to the Apoc1 cluster **C.** HCR FISH labeling of wholemount retinas, viewed en face. *Apoc1* is expressed by a subset of *C1qa^+^* microglia at P10 but not at P17. The P10 image shows six *C1qa^+^ Apoc1^+^*double-labeled cells (white arrowheads). Two examples of *C1qa^+^*cells without *Apoc1* are marked at each age (magenta arrowheads). Images are representative of staining from 4 different animals. **D.** Triple FISH labeling for *Spp1*, *Apoc1*, and *C1qa*. High-magnification view of P10 wholemount retina showing representative examples of *Apoc1^+^* microglia corresponding to the Spp1 and Apoc1 clusters, as marked. Note weaker *Apoc1* expression by the *Spp1^+^* microglial cell. Asterisk, *C1qa*-negative retinal ganglion cell expressing *Spp1*. Scale bars: 20 µm.

To verify that *Apoc1^+^* microglia are indeed present during development of the wild-type mouse retina, we performed HCR fluorescent in situ hybridization (FISH). Wholemount retinal preparations were co-stained for *Apoc1* RNA and a pan-microglial marker, *C1qa*. As predicted by the sequencing data, *Apoc1^+^*microglia were detected in situ at P5 and P10, but not at P17 (Fig. 4C). We next performed triple FISH labeling with a third marker, *Spp1*, because the Spp1 cluster also contained a large fraction of cells expressing moderate levels of *Apoc1* (Fig. 4B). Therefore, inclusion of *Spp1* probes allowed us to distinguish *Apoc1^+^*microglia originating from the Spp1 cluster (i.e., *Apoc1^+^ Spp1^+^* cells) from those representing the Apoc1 cluster (i.e., *Apoc1^+^ Spp1^−^* cells). This staining confirmed the presence of *Apoc1^+^* microglia both with and without *Spp1* (Fig. 4D). Notably, the microglia without *Spp1* tended to stain more strongly for *Apoc1* RNA than those with *Spp1* (Fig. 4C,D). This finding is consistent with the scRNA-seq data, which show that *Apoc1* is typically expressed more strongly by Apoc1 cluster cells than by microglia from the Spp1 cluster (Fig. 4B). Together, these data indicate that microglia corresponding to the Apoc1 cluster do indeed exist in vivo within developing retina.

<u>Arg1 cluster</u>. The Arg1 cluster (Supplemental Fig. S3C) was the smallest of the 16 clusters (n = 35 cells) and most selectively expressed: It was present only in the P5 dataset, and it was selectively derived from males, based on low expression of the female-specific *Xist* transcript and high expression of Y-chromosome genes within Arg1 cluster cells Supplemental Fig. S3E). Top markers of this cluster included many genes from the phagocytic expression module, including *Lgals3*, *Igf1*, and *Clec7a*, as well as several genes unique to this cluster (Supplemental Fig. S3D). This transcriptional profile is similar to an Arg1^+^ microglial population that has been identified in developing mouse brain (Hammond et al., 2019; Stratoulias et al., 2023). As in retina, the brain Arg1^+^ population was only detected during development and was extremely rare (Hammond et al., 2019), although in certain brain regions Arg1^+^ microglia are more numerous and are detectable until early adulthood (Stratoulias et al., 2023). A key difference between brain and retina relates to sex differences: Brain Arg1^+^ microglia were reported to be enriched in females rather than males (Hammond et al., 2019), or to be present at similar numbers in both sexes (Stratoulias et al., 2023). We attempted to validate the presence of Arg1 microglia in retina and to investigate potential sex differences, using FISH and using previously-validated antibodies (Stratoulias et al., 2023). However, we were unable to confidently detect Arg1^+^ microglia in P5 retinas of either sex. Given that these cells are known to exist in brain (Stratoulias et al., 2023), and given their extreme rarity within our dataset (< 1% of P5 microglia, n = 35/3837), it is possible that we simply failed to detect them histologically.

<u>Fabp5, Spp1, and Hmox1 clusters</u>. Previous scRNA-seq studies in brain identified three development-specific microglia clusters, which were named Fabp5, Spp1, and Hmox1 after marker genes enriched in each cluster (Hammond et al., 2019). We also detected corresponding clusters in our retinal dataset, based on the strong similarity of the gene sets enriched in each brain and retinal cluster (Fig. 5A,B; Supplemental Tables S1,S3). We therefore named our retinal clusters using the brain nomenclature. The Fabp5, Spp1, and Hmox1 clusters shared substantial gene expression. This was evident first from their shared expression of the phagocytic gene module (Fig. 3B). Indeed, the top marker genes of the Fabp5 and Spp1 clusters, relative to the full dataset, were mainly the core genes of the phagocytic module (Supplemental Tables S2, S3). To identify genes that distinguish these clusters from each other, we performed pairwise differential expression comparisons among the three groups (Fig. 5C). Comparing Fabp5 and Spp1, only 12 genes were significantly different between the groups, highlighting their overall similarity. Most of these (n = 10/12) were upregulated in the Spp1 state; and the two genes that were Fabp5-upregulated in this comparison (*Cd300lf* and *Slc26a11*) were not upregulated when comparing Fabp5 to Hmox1. Thus, the Fabp5 state lacks a definitive marker that can distinguish it from the other two. *Spp1* itself was by far the most differentially expressed marker distinguishing the Spp1 cluster from Fabp5 and Hmox1 microglia (Fig. 5C).

**Figure 5.**
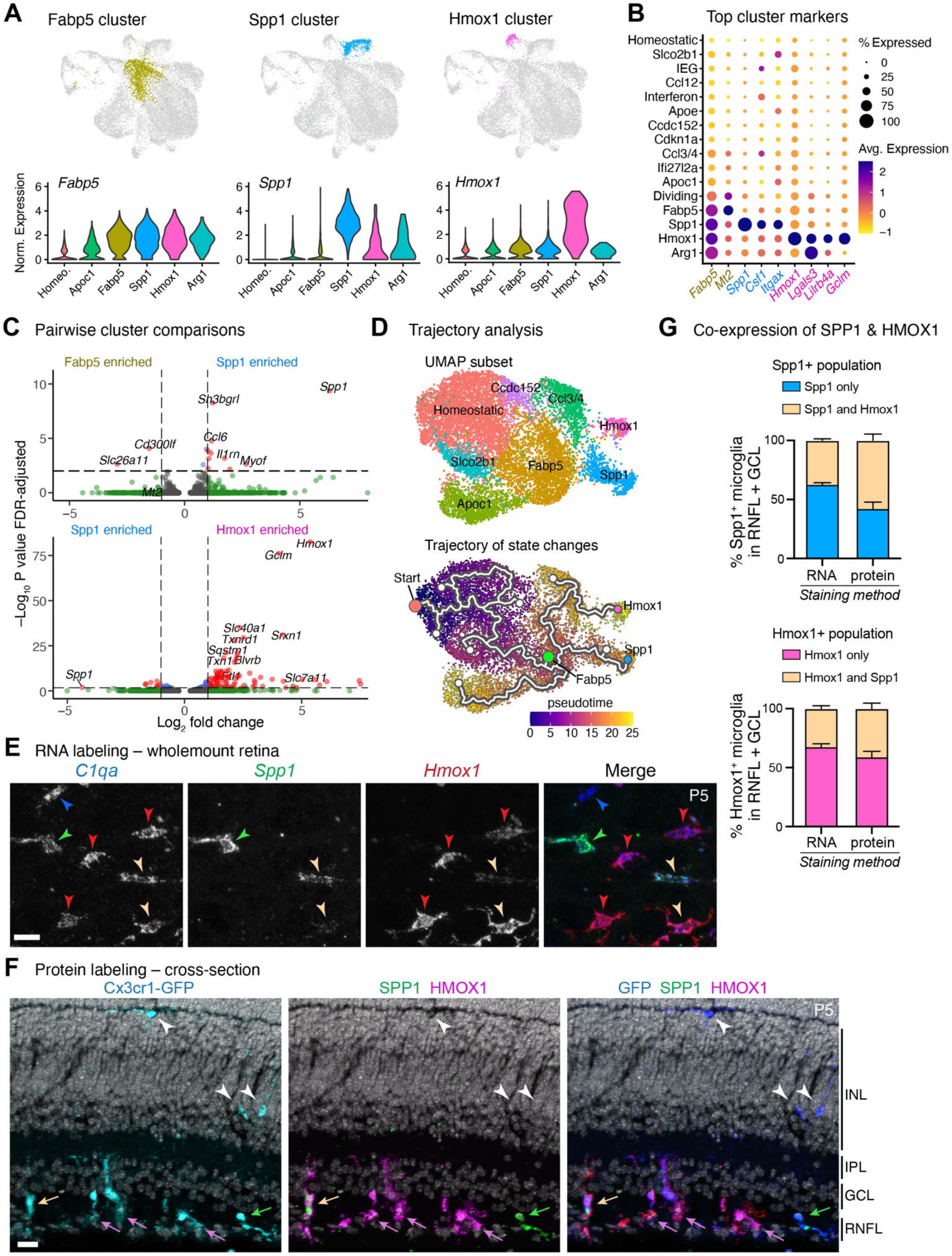
Three closely related phagocytic microglia states: Fabp5, Spp1, and Hmox1. **A.** Top row, phagocytic clusters Fabp5, Spp1, and Hmox1 highlighted on microglia UMAP plots. Bottom row, violin plots showing expression of *Fabp5*, *Spp1*, and *Hmox1* across the 5 development-specific clusters and the homeostatic cluster. **B.** Dot plot showing selected top marker genes for each cluster. **C.** Volcano plots showing pairwise gene expression comparisons between specified clusters. Red, differentially expressed genes (DEGs) that exceed P-value and fold-change cutoffs (dashed lines). Green/blue, genes exceeding only one of the cutoffs. Top: Fabp5 vs. Spp1 comparison identifies only 12 DEGs – 10 upregulated in Spp1 state and 2 upregulated in Fabp5 state. Gene labels denote top 7 DEGs. Bottom: Spp1 vs. Hmox1 comparison. Few DEGs distinguish the Spp1 state – mainly *Spp1* itself – but many DEGs are upregulated in Hmox1 microglia, including 21 known direct targets of NRF2 transcription factor (gene labels; for clarity, only 10 NRF2 targets are labeled). **D.** Trajectory inference of microglia state changes. Top, UMAP plot used for this analysis. Bottom, the same UMAP with cells colored by pseudotime, calculated relative to the homeostatic state which was defined as pseudotime = 0. Lines denote state transition paths. The analysis predicts that cells transitioning from homeostatic to phagocytic will arrive first in the Fabp5 state, before adopting Spp1 or Hmox1 phenotypes. Note that pseudotime analysis is non-directional: cells may move along the trajectory in either direction. Thus, cells may move back and forth between Spp1 and Hmox1 states, via Fabp5, by altering expression of DEGs shown in C. **E.** FISH triple-labelling for *C1qa*, *Spp1*, and *Hmox1* in P5 wholemount retina. Four distinct transcriptional states of *C1qa^+^* microglia were observed: 1) *Spp1*^−^ H*mox*1^−^ 2) *Spp1^+^*, 3) *Hmox1^+^*, 4) *Spp1^+^ Hmox1^+^* double-positive. Images depict the same representative field of view at RNFL-GCL level of a confocal Z-stack acquired from mid-peripheral retina. **F.** Immunostaining of P5 retinal cross-sections for SPP1, HMOX1, and pan-microglia marker Cx3cr1-GFP. Microglia expressing SPP1 (green arrows), HMOX1 (magenta arrows), or both (tan arrows) are localized exclusively to the RNFL-GCL region. Most such microglia have cell bodies within the RNFL; those with cell bodies in GCL still send processes into RNFL. GFP^+^ microglia within other layers lack SPP1 or HMOX1 immunoreactivity (arrowheads). **G.** The fraction of P5 RNFL microglia singly or doubly labeled for Spp1 and/or Hmox1, quantified from wholemount FISH-labeled samples similar to E (RNA), and from wholemount immunolabeled samples stained as in F (protein). Despite transcriptomic prediction that these markers should be mutually exclusive (C, bottom), double-labeled cells are common. A higher fraction of double-positive cells were detected using immunolabeling than with RNA labeling. Sample sizes: n = 4 (RNA); n = 5 (protein). Scale bars: 20 µm.

The Hmox1 group expressed many unique marker genes (Fig. 5C), but we noticed that a large fraction of these genes shared a key feature in common: they are targets of the transcription factor NRF2, a master regulator of cellular antioxidant responses. NRF2 is a ubiquitously expressed protein that only becomes transcriptionally active under conditions of oxidative stress. Using prior studies, we curated a list of 35 known and validated direct NRF2 target genes (Baird and Yamamoto, 2020; Tonelli et al., 2018); 21 of these direct targets were upregulated in the Hmox1 cluster relative to Spp1 (Fig. 5C). In addition to this stringently curated list, we also noted other likely NRF2 targets involved in antioxidant responses or iron metabolism on the Hmox1-enriched gene list (Supplemental Table S3). Thus, activity of a single transcription factor may suffice to distinguish the Hmox1 state from the Fabp5 and Spp1 states. Together, these observations suggest that all three clusters are quite similar, as they express genes that support a phagocytic function, but the Hmox1 cluster is distinguished by additional activity of the NRF2 stress-response pathway.

To validate that these microglial states exist in the retina in vivo, we performed HCR FISH staining and immunolabeling. Due to the lack of a definitive marker for the Fabp5 state, our analysis focused on the Spp1 and Hmox1 clusters. Wholemount FISH at P5 revealed that both *Spp1* and *Hmox1* RNAs are expressed by a subset of *C1qa^+^* microglia (Fig. 5E,G). Indeed, expression of both markers at this age was exclusive to microglia. By P17, however, microglia expressing these markers were not readily detectable (Supplemental Fig. S4). Thus, as predicted by the scRNA-seq data, Spp1 and Hmox1 microglial states are specific to developing retina.

Most of the *Spp1-* and *Hmox1-*positive cells in the P5 wholemount samples were detected within the portion of the confocal Z-stacks encompassing the superficial layers of the retina, such as the RNFL and GCL. This observation suggests that the Spp1 and Hmox1 populations might occupy specific retinal layers. To better evaluate the laminar location of Spp1 and Hmox1 microglia, we stained retinal cross-sections with antibodies to SPP1 and HMOX1 (Fig. 5F). Expression of both markers was restricted to microglia that were partially or fully located within the RNFL. By contrast, microglia in other layers did not express SPP1 or HMOX1 (Fig. 5F). These findings demonstrate that *Spp1* and *Hmox1* are markers of microglial populations that exist selectively in the developing retina, and that localize with striking laminar specificity to the RNFL.

### RNFL microglia switch between Spp1 and Hmox1 states

Even though the scRNA-seq data predicted that *Spp1* and *Hmox1* should be expressed by distinct microglial subsets (Fig. 5A-C), our tissue staining revealed that a substantial number of RNFL microglia expressed both markers (Fig. 5E-G). When staining for RNA, the majority of labeled microglial cells expressed only *Spp1* or *Hmox1*, confirming the separate transcriptional states predicted by the sequencing results. However, approximately one-third of the cells that expressed one of the two markers also expressed the other (Fig. 5G). One possible explanation for the presence of double positive cells is that they represent microglia in the process of downregulating one of the two markers and upregulating the other as they transition between the Spp1 and Hmox1 states. Consistent with this idea, double RNA-positive cells in many cases expressed one of the two markers much more strongly than the other (Fig. 5E). Co-staining for SPP1 and HMOX1 proteins revealed an even higher fraction of double-positive cells than the RNA staining (Fig. 5G; Fig. 6C). Given that proteins often perdure beyond the time when a cell ceases RNA expression, this result is expected under a scenario where cells are switching between two distinct transcriptional states. Thus, the pattern of RNA and protein expression suggests that the same RNFL microglial cells have a history of assuming both Spp1 and Hmox1 states.

**Figure 6.**
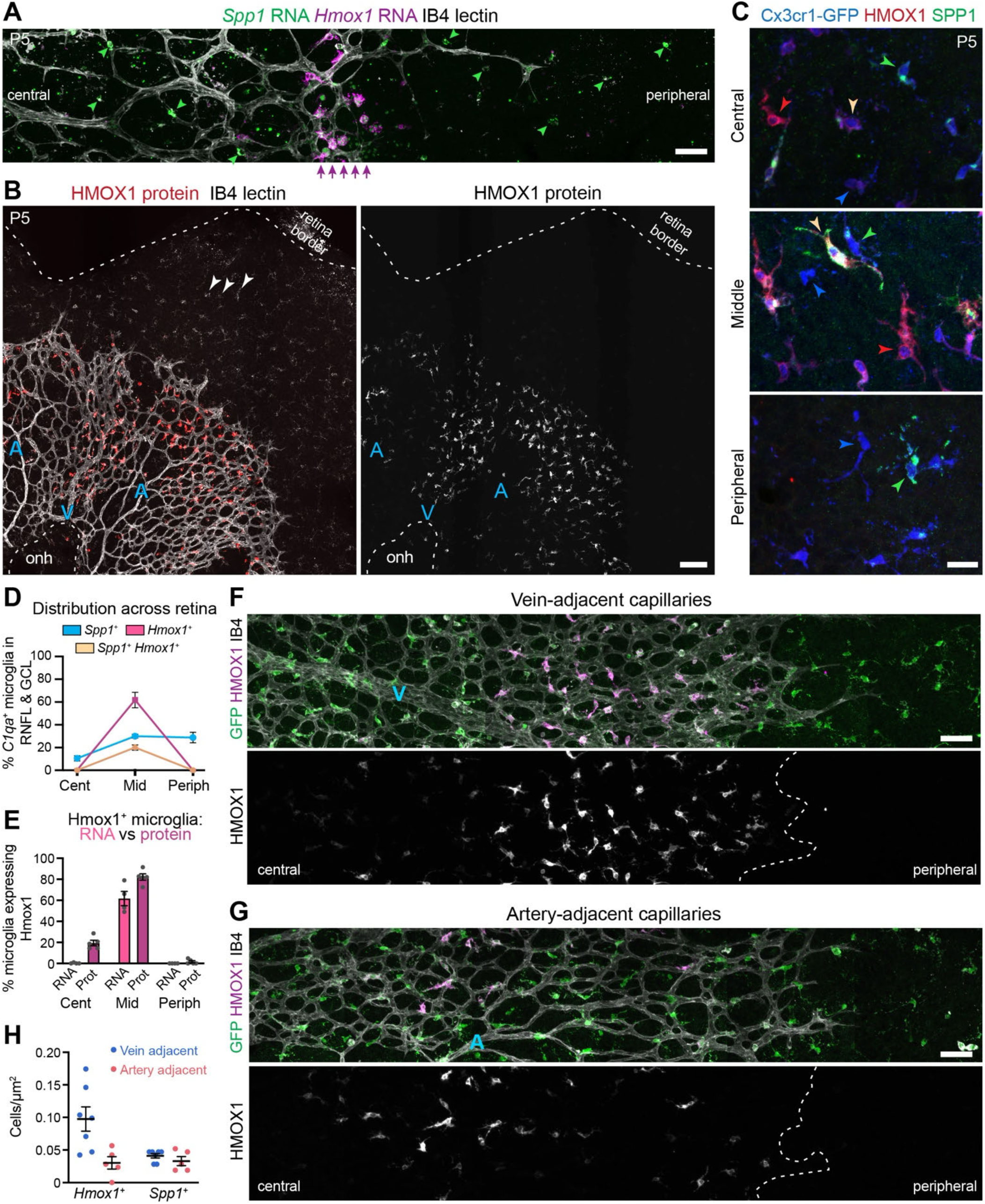
Microglial state changes associated with the angiogenic wavefront. **A.** P5 wholemount retina triple-labeled for *Spp1* and *Hmox1* RNA via FISH, and for vasculature using IB4 lectin, imaged at RNFL level. Microglia highly expressing *Hmox1* (purple arrows) are selectively located at the developing vascular wavefront, at the first vascular loops behind sprouting tip cells. *Spp1^+^* microglia (green arrowheads) are detected throughout the RNFL, on both sides of the vascular wavefront margin. **B.** Distribution of HMOX1 protein-expressing microglia across P5 retina. Left, representative stitched image of RNFL from a retinal wholemount double stained for HMOX1 and IB4 lectin, which labels vasculature and microglia (arrowheads). Right, same field of view showing only HMOX1 channel. Dashed line, edge of retina. Microglia ahead of the wavefront do not express HMOX1 protein, but microglia at the wavefront are strongly labeled. Behind the wavefront, microglia with high HMOX1 protein labeling extend further into central retina as compared to *Hmox1* RNA (A), especially in regions adjacent to veins (‘V’). Fewer HMOX1^+^ microglia are found near arteries (‘A’). ONH, optic nerve head. **C.** Expression of SPP1 and HMOX1 protein by Cx3cr1-GFP^+^ microglia in central, mid-peripheral (middle), and peripheral P5 retina. Both SPP1 and HMOX1 are exclusively expressed by GFP^+^ microglia at this age. Numerals, examples of GFP^+^ microglia expressing different marker combinations: 1) Negative for both SPP1 and HMOX1; 2) SPP1*^+^*only; 3) HMOX1*^+^* only; 4) double-positive for SPP1 and HMOX1. **D.** Prevalence of P5 RNFL/GCL microglia expressing *Spp1* and/or *Hmox1* RNA across different eccentricities, quantified from wholemount FISH images similar to Fig. 6E. *Spp1^+^* microglia are found throughout RNFL, but *Hmox1*^+^ microglia are exclusively found in middle retina, where the vascular wavefront is located at this age (see A,B). Some *Hmox1^+^* microglia co-express *Spp1*, suggesting that cells in the Spp1 state can activate *Hmox1* RNA expression upon arrival of vasculature. Sample size: n = 4 mice. **E.** The fraction of RNFL microglia expressing *Hmox1* RNA (same data as D) compared to the fraction expressing HMOX1 protein (quantified from images similar to C). HMOX1 protein expression persists in microglia of central retina, behind the vascular wavefront, that, no longer express *Hmox1* RNA. Sample size (protein): n = 5 mice. **F,G**. Representative en face images of the vascular wavefront region in P5 wholemount retinas, showing distribution of HMOX1^+^ microglia near veins (‘V’) or arteries (‘A’). More of the Iba1^+^ microglia population co-expresses HMOX1 in vein-adjacent capillary beds (F) than in artery-adjacent capillaries (G). Dashed lines, vascular wavefront. **H.** Quantification of microglia expressing *Spp1* or *Hmox1* RNA in wavefront regions adjacent to veins or arteries. Significantly more *Hmox1*^+^ cells were found in vein regions (P = 0.011; T-test corrected for multiple comparisons) Sample sizes: n = 7 veins and n = 5 arteries from 2 mice. Scale bars: 50 µm (A, F, G); 100 µm (B); 20 µm (C).

To learn whether the scRNA-seq dataset supports this model, we performed trajectory analysis (Trapnell et al., 2014). This method uses graded changes in gene expression to order cells along a pseudotime axis, enabling predictions as to how cells transition between states. Trajectory inference was performed on a filtered dataset in which small/minor clusters and the artifactual IEG cluster were removed, thereby focusing the analysis on major clusters of interest (Fig. 5D, top). Importantly, pseudotemporal ordering is inherently non-directional, meaning that cells could in principle move in either direction along the inferred trajectories. Here we defined the homeostatic state as the trajectory’s starting point – i.e. the state with a pseudotime value of zero – but this choice was arbitrary and does not affect the shape of the trajectory (Fig. 5D, bottom).

The inferred state-change trajectory suggests the Fabp5 cluster defines a baseline “hub” state, from which microglia can move into multiple non-homeostatic states including Spp1 or Hmox1 (Fig. 5D). The notion that Fabp5 is a baseline state is consistent with our failure to find unique markers that distinguish it from the other phagocytic states (Fig. 5C). According to the inferred trajectory, cells transitioning between Spp1 and Hmox1 states do so by passing through the baseline Fabp5 state. Differential gene expression analysis suggests that such a transition could be accomplished by changing expression of only a few genes. To move from Spp1 to Fabp5, the main gene needing downregulation is *Spp1* itself (Fig. 5C). Activating NRF2 transcription would then suffice to drive the transition to Hmox1 (Fig. 5C). The reverse state change could be accomplished by turning off NRF2-driven transcription and upregulating *Spp1*. Together, these analyses show that it is plausible for microglial cells to move rapidly between the Spp1 and Hmox1 states, thereby supporting the view that *Spp1 Hmox1* double-labeled cells detected histologically are indeed undertaking this state transition in vivo.

### State changes of RNFL microglia are associated with developing vasculature

We next sought to understand the developmental events associated with switching between Spp1 and Hmox1 states. To this end, we determined the retinal location of RNFL microglia expressing *Spp1* and/or *Hmox1*. Using both antibody and RNA labeling on wholemount P5 retina, we found that microglia highly expressing *Hmox1* were exclusively located in central and mid-peripheral retinal regions, but were excluded from the far periphery (Fig. 6A-D). By contrast, *Spp1^+^* microglia were distributed uniformly across retinal eccentricities (Fig. A,C,D).

One of the key differences between central and peripheral retina at P5 is that the periphery has yet to become vascularized. Retinal angiogenesis sweeps through the RNFL in a wave-like manner during the first postnatal week, spreading outward from the optic nerve head to reach the periphery by P7-8 (Paisley and Kay, 2021). At P5, the wavefront of the developing vasculature is located in middle retina, raising the possibility that the Hmox1 state could be associated with arrival of vessels. Co-staining for vascular markers revealed that *Hmox1* RNA^+^ microglia were strictly localized to the advancing angiogenic wavefront, aligning with remarkable precision to the first vascular loops behind the immature tip cells at the vanguard (Fig. 6A,D). HMOX1 protein showed a similar expression pattern, except that strong immunoreactivity was also detected in RNFL microglia further behind the wavefront, extending into central retina (Fig. 6B,C,E).

The central-most microglia were often more weakly immunoreactive for HMOX1 than those at the vascular wavefront (Fig. 6B,F), consistent with their lack of active *Hmox1* RNA expression (Fig. 6A,E). Together, this RNA and protein expression pattern suggests that arrival of vessels at a given retinal location triggers a transient pulse of *Hmox1* RNA and protein expression, followed by gradual protein degradation as the wavefront advances. A notable fraction of the newly *Hmox1*^+^ microglia at the vascular wavefront co-expressed *Spp1* RNA (Fig. 6D), suggesting that they were in the process of transitioning between states. This observation is consistent with a model in which at least some of the RNFL microglia in the Spp1 state are induced to switch into the Hmox1 state upon vessel arrival.

Further analysis of HMOX1-immunoreactive microglia revealed additional specificity in their localization relative to developing vasculature. Initially, we noticed that some regions of central retina contained far more HMOX1^high^ microglia than other adjacent regions. Co-staining for vasculature revealed that this pattern was associated with the location of retinal arteries and veins (Fig. 6B). Even near to the vascular wavefront, RNFL microglia in the Hmox1 state were far more numerous in capillary beds adjacent to developing veins than those adjacent to arteries (Fig. 6F,G). This finding suggests that there may be additional vessel type-specific factors determining RNFL microglia state.

### Phagocytic function of Spp1 and Hmox1 RNFL microglia

Our module score analysis (Fig. 3B) indicates that Spp1 and Hmox1 microglia are likely to engage in phagocytic functions. If this is true, Spp1 and Hmox1 microglia should exhibit anatomical features indicative of phagocytic activity (Ransohoff and Perry, 2009). We therefore performed anatomical studies using antibodies to SPP1 and HMOX1, in which we tested for three specific hallmarks of phagocytic function: 1) A simplified, unbranched arbor morphology, as opposed to the highly ramified morphology that typifies homeostatic microglia; 2) Extension of phagocytic cups that engulf extracellular material; and 3) The presence of intracellular compartments/organelles containing foreign material. We limited our analysis to cells highly expressing the two protein markers (SPP1^high^ and/or HMOX1^high^), so as to maximize the chances that the analyzed cells also express the corresponding RNA (Fig. 6E) and therefore correspond to the Spp1 or Hmox1 transcriptional states.

RNFL microglia expressing these two markers, either singly or jointly, consistently exhibited a simple amoeboid morphology without extensive arbor ramification (Fig. 7B-D). Further supporting their classification as phagocytic microglia, phagocytic cups were frequently observed extending from these SPP1^high^ and/or HMOX1^high^ cells, and intracellular compartments containing phagocytosed material were often present (Fig. 7B-F). Some *Hmox1* RNA^+^ microglia at the vascular wavefront, especially in venous regions, showed a particularly extreme version of this morphology, with a highly rounded arbor-free morphology and a cytoplasm that was filled with multiple vacuole-like structures (Fig. 7G). These numerous internal compartments were in many cases filled with autofluorescent material (Fig. 7G) – possibly lipofuscin (Stillman et al., 2023), or possibly heme derived from engulfed red blood cells.

**Figure 7.**
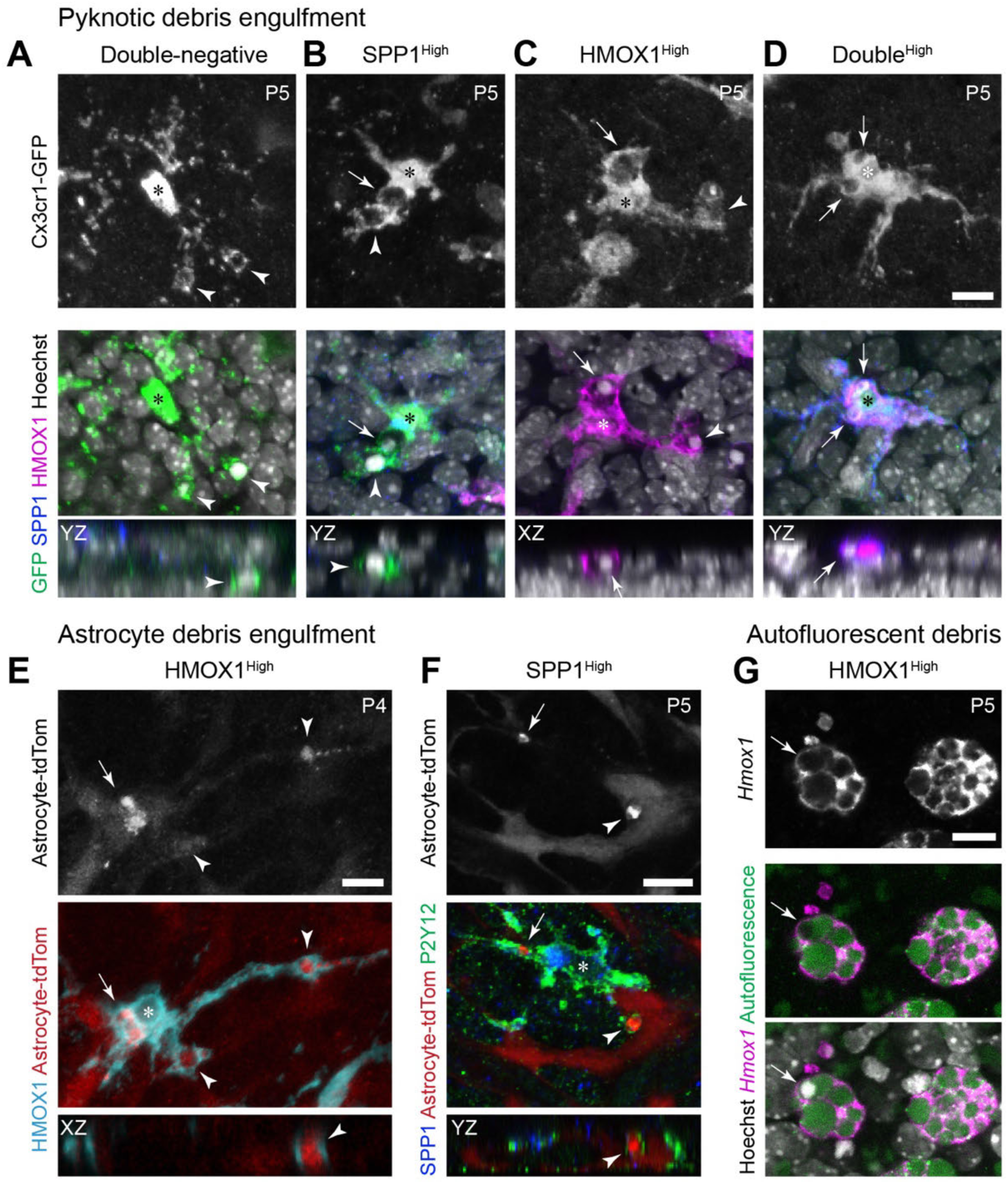
Spp1 and Hmox1 microglia engulf cellular debris during normal development. **A-D**. Phagocytic morphology of microglia in P5 retina. Top row, en face Cx3cr1-GFP images of microglia from mid-peripheral retina, showing representative morphologies of cells with the specified marker expression profile. A: Microglial cell lacking SPP1 and HMOX1, from inner nuclear layer. Note ramified morphology. B-D: SPP1- and/or HMOX1-expressing microglia in RNFL exhibit amoeboid morphology. Middle and bottom rows, merged multi-channel images of same cells, viewed en face (middle) or in XY or YZ orthogonal projections (bottom), showing expression of specified markers (GFP channel was omitted in C,D for clarity; GFP was redundant because HMOX1 shows cell morphology). SPP1- and/or HMOX1-positive microglia (B-D) have phagocytic cups (arrowheads) surrounding pyknotic nuclei (small bright Hoechst-positive bodies); and they contain internalized pyknotic nuclei within cytoplasmic organelles (arrows). Double-negative microglia (A) can still have phagocytic cups, but rarely exhibit cytoplasmic inclusions containing pyknotic debris. **E,F**. Astrocyte debris engulfment by Hmox1 (E) and Spp1 (F) microglia. Top row, astrocytes transgenically labeled with tdTomato (tdTom) reporter. Astrocyte debris (arrows, arrowheads) is punctate and brighter than the cytosolic tdTom of live astrocytes. Middle and bottom rows, merged multichannel images, en-face (middle) or orthogonal (bottom) views, showing labeling for specified markers. Microglia morphology is revealed by anti-HMOX1 (E) or anti-P2Y12 (F). Astrocyte debris localizes to phagocytic cups (arrowheads) or somatic compartments (arrows) of both Hmox1 and Spp1 microglia. **G**. A subset of Hmox1 microglia entirely lacked arbors and were packed with vacuole-like intracellular compartments containing autofluorescent material. Arrow, internal compartment containing a pyknotic nucleus; this compartment was not autofluorescent, indicating that autofluorescent material is not comprised of apoptotic cell debris. Scale bars, 10 µm; bar in (D) applies to (A-C).

To learn whether these anatomical features are specific to SPP1^high^ and/or HMOX1^high^ microglia, we compared them to double-negative cells, identified from P5 retina within deeper retinal layers or from mature P17 retina (Fig. 7A; Supplemental Fig. S4). While P5 double-negative cells still typically had phagocytic cups, consistent with the paucity of homeostatic microglia at this age (Fig. 3C), their morphology was typically far more ramified and complex than the amoeboid SPP1^high^ or HMOX1^high^ microglia (Fig. 7A-D). Vesicles containing debris were not observed within the somata of double-negative cells (Fig. 7A; Supplemental Fig. S4). Together these observations suggest that, while they are not the only phagocytic cells in P5 retina, SPP1^high^ and/or HMOX1^high^ microglia have anatomical hallmarks of cells that are particularly phagocytic.

We next investigated what kind of material SPP1^high^ and HMOX1^high^ microglia might be engulfing. During the first week of postnatal mouse development, microglia of the inner retina have two main phagocytic targets: 1) neurons undergoing naturally-occurring developmental death via apoptosis; and 2) astrocytes undergoing non-apoptotic developmental death mediated by microglia (Anderson et al., 2019; Puñal et al., 2019). To assess engulfment of apoptotic neurons, we used Hoechst DNA dye to identify pyknotic nuclei, a reliable anatomical correlate of neuronal apoptosis. Apoptotic bodies were readily identified within SPP1^high^ and HMOX1^high^ microglia, either encircled by phagocytic cups or contained within somatic vacuole-like compartments (Fig. 7A-D). To assess engulfment of astrocytes, we used an astrocyte-specific tdTomato transgenic marker and surveyed microglia for tdTomato-positive debris. Both SPP1^high^ and HMOX1^high^ microglia were found to engulf astrocyte material, which again could be observed both within phagocytic cups and within intracellular compartments (Fig. 7E,F). These results indicate that both Spp1 and Hmox1 microglial states are associated with clearance of cellular debris generated during the normal course of postnatal retinal development.

## DISCUSSION

In this study we have performed a comprehensive survey of microglia and macrophage diversity across postnatal development of mouse retina. We show that transcriptional diversity is driven in many cases by the precise location of myeloid cells within the developing eye. In the case of macrophages, BAM subtypes bordering the inner and outer retina express distinct molecular phenotypes: Lyve1 BAMs localize to vitreous and lens (i.e. inner borders), while MHCII BAMs localize to RPE and choroid (i.e. outer borders). And in the case of microglia, the phagocytic Spp1 and Hmox1 states localize selectively to a single retinal layer, the RNFL. While developing microglia in other layers exhibit phagocytic activity (Jiang et al., 2022), our data suggest such microglia rarely assume either of these two states – at least at the ages studied here.

Furthermore, the Hmox1 state showed additional anatomical specificity in its selective association with angiogenic sprouting at the vascular wavefront. Unlike the previously-described ‘capillary-associated microglia’ state, which is stably expressed by vasculature-adjacent microglia in mature brain (Bisht et al., 2021; Mondo et al., 2020), the Hmox1 state was expressed only transiently during development in a manner that correlated with the arrival of endothelial tip cells and/or the onset of circulation. These anatomical results highlight how our scRNA-seq dataset can provide a useful framework for interrogating how distinct myeloid populations support diverse developmental functions at diverse ocular locations.

### Comparison with previous scRNA-seq studies of CNS myeloid cells

Our sequencing results are broadly in line with a previous scRNA-seq study of developing retinal microglia at a single age (P6-P7) (Anderson et al., 2022). However, the present study extends our knowledge of retinal myeloid diversity in several important new ways. First, because our dataset covers a broader range of ages, we were able to document how microglia diversity changes over time. This information, together with our multiple approaches to pinpointing ex vivo transcriptional artifacts, has enabled definitive identification of development-specific transcriptional states that are authentic to retinal microglia in vivo. Second, our dataset also includes macrophages, allowing us to identify the ocular location of BAM populations that were previously described in brain (Utz et al., 2020; Van Hove et al., 2019). Finally, our dataset contains a far larger sample size of microglia from wild-type mice, providing greater resolution to detect distinct transcriptional states via unsupervised clustering. This advantage revealed previously unappreciated diversity within known transcriptional types. For example, Anderson et al. (2022) identified a large “Apoe-enriched” cluster, which was thought to be a single population devoted to phagocytosis and tissue remodeling.

However, our data suggest that this large cluster likely comprises multiple distinct populations, including the phagocytic Fabp5 cluster, but also the homeostatic Apoe and Ccdc152 clusters which are less likely to engage in tissue remodeling based on their module scores (Fig. 3B). This distinction, enabled by our new data, helps to further narrow down the molecular phenotype of the microglia that engulf apoptotic neurons and that are diminished when neuronal apoptosis is blocked (Anderson et al., 2022).

Comparing mouse retina and brain scRNA-seq data, we can draw several conclusions about myeloid cell commonalities across these CNS regions. First, in both regions, microglia are far more transcriptionally heterogeneous during development than at maturity (Hammond et al., 2019; Li et al., 2019), suggesting that microglia undertake a range of specialized developmental functions that are no longer needed in adulthood. Macrophages, by contrast, do not exhibit development-specific states in brain (Li et al., 2019) or retina, suggesting that BAMs are unlikely to engage in development-specific functions requiring transcriptional specialization. Second, the particular transcriptional states observed for developing myeloid cells in each CNS region are broadly similar – especially in the healthy/uninjured setting. Microglial states found in both healthy mouse brain and retina include the Homeostatic, Interferon, and Ccl3/4 clusters, as well as the four phagocytic clusters – Arg1, Fabp5, Spp1, and Hmox1 (Escoubas et al., 2024; Hammond et al., 2019; Marsh et al., 2022; Stratoulias et al., 2023). The Lyve1 and MHCII BAM states were also found in both retina and brain under homeostatic conditions (Brioschi et al., 2023).

Despite these similarities, there were some differences uncovered by our comparisons of myeloid scRNA-seq studies from mouse retina and brain. First, there was a pair of microglial states – Ifi27l2a and Apoc1 – that were prevalent in our retinal dataset but were rare in healthy brain, instead becoming upregulated by brain microglia mainly in injury or disease contexts (Hammond et al., 2019; Masuda et al., 2019; Somebang et al., 2021). Another group of clusters from our dataset – Slco2b1, Ccl12, and Cdkn1a – did not have a clear match to previously reported scRNA-seq clusters from brain, even though these three top marker genes are known to be expressed by brain microglia in certain contexts. *Slco2b1* encodes a heme transporter that is broadly expressed by brain microglia (Unlu et al., 2022). Our data were mostly consistent with this observation, as we found *Slco2b1* to be expressed at moderate levels by all retinal microglia; however, expression was approximately twofold higher for cells of the Slco2b1 cluster. *Ccl12* is upregulated by brain microglia in injury settings, as shown by several scRNA-seq studies (Hammond et al., 2019; Mendiola et al., 2020; Witcher et al., 2021). However, the other markers of these injury-induced *Ccl12^+^* scRNA-seq clusters do not match our retinal *Ccl12* cluster, suggesting that developmental *Ccl12* expression is driven by a distinct mechanism. *Cdkn1a* is often used as a marker of senescent microglia; however, other typical senescence markers were not expressed in our retinal cluster. Further exploration of these brain-retina microglial differences will be an important direction for future studies.

Finally, there was one cluster in our retinal microglia dataset – the cluster marked by *Ccdc152* expression – that is not only absent from brain scRNA-seq datasets, but is also marked by a gene that has rarely been studied. There is minimal information available about the function or expression pattern of the *Ccdc152* gene within the CNS. To our knowledge, its expression by microglia in mouse or human has not previously been reported. Because it did not mark a development-specific population, we did not validate microglial expression of *Ccdc152* using histology. Thus, further studies will be needed to confirm existence of this microglial population.

### Diversity of phagocytic microglia in developing retina

Our data provide new insights into the transcriptional diversity and potential developmental functions of the phagocytic microglia populations. Microglia corresponding to our Fabp5 and Spp1 clusters have previously been characterized in both retina and brain (Anderson et al., 2022; Ghena et al., 2025; Hammond et al., 2019; Li et al., 2019). These microglia express a well-characterized gene signature, described under many different names (e.g. PAM, ATM, DAM), that is found exclusively in settings where microglia engage in substantial phagocytic activity – such as development or certain disease contexts (for review see (Anderson and Vetter, 2019; Benmamar-Badel et al., 2020). During development, microglia with this gene profile engulf apoptotic neurons; and in the neonatal brain they localize preferentially to developing axon tracts where they engulf excess oligodendrocyte precursors (Anderson et al., 2022; Barclay et al., 2024; Hagemeyer et al., 2017; Hammond et al., 2019; Nemes-Baran et al., 2020; Wlodarczyk et al., 2017). Here we find that retinal Spp1 microglia also localize selectively to an axon tract layer – i.e., the RNFL. Even though the retina lacks oligodendrocytes, it is plausible that phagocytic load would be greater in the RNFL than in other retinal regions, because, while neurons undergo apoptosis in all layers, only the RNFL contains astrocytes which die in massive numbers during the first two postnatal weeks (Puñal et al., 2019). This excess phagocytic burden may explain the paucity of Spp1 microglia in other layers.

We also identify for the first time that Hmox1 microglia represent a phagocytic state with high similarity to the Fabp5 and Spp1 states. Hmox1 or antioxidant-expressing microglial clusters have been noted in earlier scRNA-seq studies (Anderson et al., 2022; Hammond et al., 2019; Mendiola et al., 2023), but to our knowledge there have been few efforts to locate such cells in developing brain or to investigate their function. Unlike Fabp5 and Spp1 microglia, the Hmox1 cells were not considered part of the phagocytic (i.e PAM-ATM-DAM) microglial population. Here we show that the phagocytic gene module is in fact shared among all three of these states, and that the major distinguishing feature of Hmox1 microglia is the additional upregulation of a suite of genes typically controlled by NRF2, the master regulator of cellular antioxidant responses. Our histological studies validate the existence and phagocytic nature of Hmox1 microglia in the retina in vivo. Moreover, we demonstrate that the three phagocytic states are closely linked, with frequent switching especially among Spp1 and Hmox1 transcriptional states. While we did not have Cre-loxP fate mapping tools available to assess state switching, we instead leveraged perdurance of SPP1 and HMOX1 protein to show that many RNFL microglia – particularly those behind the developing vascular wavefront – have a history of occupying both Spp1 and Hmox1 states.

Why would developing microglia need to switch among three different phagocytic states? To handle a heavy phagocytic load, microglia need to metabolize and degrade ingested material in preparation for ingesting more. An appealing model is that each of the closely related transcriptional states – Fabp5, Spp1, and Hmox1 – support distinct metabolic needs of phagocytic microglia. Many of the phagocytic module genes are involved in lysosomal function or lipid metabolism, suggesting that they support degradation of engulfed material. Because this gene module is particularly highly expressed in Spp1 and Hmox1 states (Fig. 3B,G), shifts into these states may reflect greater metabolic demands driven by increased phagocytic load. Shifts from the Spp1 to Hmox1 state appear to be spurred by increases in oxidative stress, based on the NRF2 gene signature that distinguishes these states. One plausible source of oxidative stress is increased mitochondrial respiratory activity – driven either by the metabolic demands of engulfment and digestion (He et al., 2021; Ishii et al., 2000), and/or by the sudden increase in oxygen availability coinciding with arrival of the angiogenic wavefront. Additionally, our observation of autofluorescent material within Hmox1 microglia indicates that metal ions may be present in their lysosomes, which can be a major source of reactive oxygen species (Gray and Woulfe, 2005; Moreno-Garcia et al., 2018). Upregulation of the Hmox1 expression profile may serve to alleviate these sources of oxidative stress, thereby returning microglia to Spp1 and/or Fabp5 states where they can continue their debris-clearing activities.

## Conclusions

Altogether, this survey of myeloid cell populations in the early postnatal mouse retina provides a framework for future studies investigating the role of these cell types in CNS development. By showing that particular microglia and macrophage transcriptional states are present at specific times and places during eye development, our study opens the way to future investigations of how those transcriptional states support developmental events that are occurring in their vicinity. While we have only performed in vivo validation for a subset of the transcriptional states predicted by the scRNA-seq analysis, the successful validations we did perform increase confidence that future studies will indeed confirm the presence of the remaining populations. The overall similarity of microglial populations within developing brain and retina suggests that our findings could have general implications for understanding microglia functions in CNS development.

## METHODS

### Data Availability

Sequencing data have been deposited at NCBI GEO – accession number GSE293912.

### Mice

Animal experiments were reviewed and approved by the Institutional Animal Care and Use Committee of Duke University. Mice of both sexes were used. The animals were maintained under a 12 h light-dark cycle with ad lib access to food and water.

The following strains were obtained from Jackson Laboratories: C57BL6/J (RRID:IMSR_JAX:000664); *Cx3cr1^CreER-ires-YFP^* (RRID:IMSR_JAX:021160); *Cx3cr1^GFP^* (RRID:IMSR_JAX:005582); *Ccr2^RFP^* (RRID:IMSR_JAX:017586); lox-stop-lox-Sun1GFP (*Gt(ROSA)26^Sortm5.1(CAG-Sun1/sfGFP)Nat^/MmbeJ*, RRID:IMSR_JAX:030952); GFAP-Cre (Tg(GFAP-cre)25Mes, RRID:IMSR_JAX:004600); Ai14 tdTomato Cre reporter (*Gt(ROSA)26Sor^tm14(CAG-tdTomato)Hze/J^*; RRID:IMSR_JAX:007914).

The Aldh1l1-tdTomato line (STOCK Tg(Aldh1l1-tdTomato)TH6Gsat/Mmucd; RRID:MMRRC_036700-UCD) was obtained from the MMRRC repository. Wild-type CD-1 mice (Crl:CD1(ICR), RRID:IMSR_CRL:022) were obtained from Charles River.

*Husbandry and breeding strategies*: All mutant and transgenic strains were maintained on the C57Bl6/J background. Strains were genotyped for background mutations affecting retinal health (*rd1*, *rd8*) and any carriers of these deleterious mutations were eliminated from the colony. To fluorescently label microglia for flow cytometry and histology, homozygous *Cx3cr1^CreER-ires-YFP^* or *Cx3cr1^GFP^* animals were outcrossed to C57Bl6/J breeders, and YFP^+^ or GFP^+^ cells were identified by native fluorescence (FACS) or anti-GFP staining (histology). To fluorescently label myeloid cell nuclei, *Cx3cr1^CreER-ires-YFP^*mice were crossed to the lox-stop-lox-Sun1GFP line followed by administration of tamoxifen (100 µg), which resulted in Cre-dependent expression of the nuclear envelope-targeted Sun1GFP construct. To label astrocytes for anatomy experiments, two different genetic strategies were used: 1) The Ai14 tdTomato Cre reporter line was crossed to the astrocyte-selective GFAP-Cre driver line; and 2) the Aldh1l1-tdTomato line was used to directly express fluorescent protein expression within astrocytes.

### Single-cell sequencing sample preparation

*Cx3cr ^CreER-ires-YFP^* mice were anesthetized and euthanized via decapitation, then eyes were enucleated into ice cold oxygenated AMES media. Tissues and cells were kept on ice throughout the procedure except as noted, and centrifugation was carried out at 4°C. To isolate the retina, the anterior portion of the eye was removed and the retina was dissected free of the eyecup. The retinal pigment epithelium typically remained partially attached upon removal from the eyecup, and if so, no further efforts were made to peel it off. All YFP-positive mice in each litter were used – typically 5-7 animals yielding 10-14 retinas, which were pooled for downstream steps.

Retinal tissue was dissociated in a solution of activated papain (Worthington, 16 U/mL) at 37°C for 25 to 30 min, then triturated in an ovomucoid solution (Worthington, 15 mg/mL) to prepare a single cell suspension. The suspension was passed through a 30 μm cell strainer and pelleted at 400 g for 7 min. The cells were resuspended at 1×10^7^ cells/mL in AMES solution and incubated in LIVE/DEAD Fixable Aqua Dead Cell Stain (Thermo, 1:1000) for 30 min at 4°C. Cells were washed twice, centrifuged at 400 g for 2 min, and resuspended in 0.4% BSA in 1X PBS. The YFP^+^ live population was collected via flow cytometry (MoFlo Astrios or BD Diva cytometers) and sorted into 0.4% BSA in 1X PBS. Post-sorting, samples were pelleted and resuspended in PBS + 0.04% BSA, and were then processed by the Duke Molecular Genomics Core for 10x Genomics droplet encapsulation and library preparation (Chromium Single Cell 3’ assay, v3). Samples were sequenced on Illumina NextSeq or NovaSeq instruments.

Samples were collected at P5, P10, and P17, with two replicates (i.e. litters) per timepoint. Tamoxifen was not administered for these studies. One of the P5 samples did not pass sequencing quality control (see below) and was not used for further analysis. For experiments that included transcriptional/translational inhibitors: Actinomycin D (Sigma #A1410; 5µg/mL); Triptolide (Sigma #T3652; 3.7µg/mL, i.e. 10µM); and Anisomycin (Sigma #A9789; 27.1µg/mL) were added to all solutions, using working concentrations noted above as previously described (Marsh et al., 2022).

### Single-nucleus sequencing sample preparation

*Cx3cr1 ^CreER-ires-YFP^*; lox-stop-lox-Sun1GFP mice were injected with tamoxifen at P0-1 and collected at P5 or P10. One litter (5-7 mice; 10-14 retinas) was collected at each age. As in the single-cell preparation, retinas were isolated in ice-cold oxygenated AMES media. Retinas were then flash-frozen in liquid nitrogen and stored at –80°C. To prepare a nuclear suspension, frozen retinas were thawed for 5 min on ice; incubated for 5 min on ice in lysis buffer (10 mM Tris-HCl pH 7.5, 10 mM NaCl; 3 mM MgCl_2_; 10% Nonidet-P40 substitute; 40U/µL RNAse inhibitor); triturated 20 times with a wide-bore pipette tip; incubated on ice an additional 10 min; and homogenized with 20 strokes of a disposable microcentrifuge tube dounce homogenizer. The preparation was then suspended in 0.5 mL AMES + 2% BSA containing 40U/µL RNAse inhibitor; passed through at 30 µm cell strainer; and pelleted at 500 g for 5 min at 4°C before resuspending in AMES + 2% BSA. Flow cytometry isolation of GFP^+^ nuclei and downstream steps were as described in the single-cell procedures above.

### Analysis and processing of 10x datasets

For each sample, reads were demultiplexed using bcl2fastq (Illumina), and aligned to the mm10 reference genome using Cellranger count (v5.0.1). For single-nucleus but not for single-cell datasets, the Cellranger analysis counted reads that mapped to introns. The Cellranger pipeline generated raw and filtered feature-barcode matrices using default parameters. SoupX v1.6.2 (Young and Behjati, 2020) was used to estimate contamination from ambient RNA based on the raw and filtered matrices, and to generate a corrected count matrix based on these estimates. The corrected count matrix was saved in Cellranger feature-barcode output format using the DropletUtils package (v1.18.1). One of the samples, P5 replicate 2, was excluded from further analysis because there were few intact cells and high levels of ambient RNA contamination.

Seurat version 4.3.0 (Stuart et al., 2019) was used for downstream analysis steps including filtering of low quality cells, normalization, dimensionality reduction, clustering, and identification of differentially expressed genes (DEGs). Cells with low sequencing quality were identified and filtered based on several criteria, including high fraction of mitochondrial or ribosomal gene reads, or low numbers of detected genes and unique molecular identifiers (UMIs). Cutoffs for these parameters varied slightly by dataset, but in most cases we discarded cells that failed to meet the following thresholds: Less than 10% mitochondrial reads; less than 25-30% ribosomal reads; greater than 1,500-2,000 genes; greater than 2,500-3,000 UMIs.

Filtered samples were processed using the Seurat SCTransform pipeline with the mitochondrial and ribosomal percentage variables regressed from the model. Seurat::RunPCA, RunUMAP, FindNeighbors and FindClusters functions were used for dimensionality reduction (using the top 30 PCA dimensions) and unsupervised clustering. To prevent variations in the number of males or females within each sequenced litter from exerting an undue influence on clustering, we excluded any male-specific (Y chromosome) or female-specific (*Xist*, *Tsix*) transcripts from the variable features used for dimensionality reduction. An initial clustering resolution of 0.8 was used for the FindClusters function.

To integrate the 5 individual single-cell datasets and to control for batch effects, we used the Seurat Reciprocal PCA integration workflow. Each individual dataset was prepared for integration by first removing contaminating neurons and astroglia. These contaminants were identified by low expression of the myeloid marker *Cx3cr1*, and high expression of cell type specific markers (neurons: *Nrxn3, Rho, Isl1, Tfap2b*; astroglia: *Sox9*). To generate the microglia-only integrated object (Fig. 3), we additionally filtered the individual datasets to remove macrophages and monocytes, identified by low expression of the microglia marker *Sparc* and high expression of macrophage markers (*Mrc1, Ms4a7, Pf4*) or monocyte markers (*Ccr2*, *Plac8*). Filtered datasets were integrated using the following Seurat functions (key arguments noted in parentheses): 1) SelectIntegrationFeatures (nfeatures = 3000); 2) PrepSCTIntegration; 3) FindIntegrationAnchors (normalization method = SCT, dims = 1:30, reduction = “rpca”, k.anchor = 20); 4) IntegrateData (normalization method = SCT; dims = 1:30). Integrated objects underwent dimensionality reduction and clustering as described above, using the Seurat ‘Integrated’ assay as the data source. To visualize the single-nucleus data, we used the Seurat::MapQuery function to project nuclei onto the UMAP coordinates defined by the main integrated single-cell dataset.

### Differential gene expression analysis

DEGs for each cluster were identified using the Seurat::FindAllMarkers function using the default Wilcoxon Rank Sum test, which was run on the normalized data from the RNA assay (SCT and Integrated assays were only used for dimensionality reduction and clustering). Feature plots mapping gene expression levels across cells, and dot plots comparing gene expression across clusters, were made using the scCustomize package (Marsh, 2021). Violin plots were produced with the Seurat::VlnPlot function.

For volcano plots in Fig. 2 and Fig. 5, we performed a pseudobulk analysis by averaging gene expression across clusters within individual datasets. For the phagocytic cluster comparisons, three cell samples (P5-replicate1, P10-replicate1, P10-replicate2) were compared using the DESeq2 package. For the cell vs. nucleus comparisons, the same three cell samples were compared to two nucleus samples (P5 and P10) using DESeq2. Data were plotted with the EnhancedVolcano package.

### Module score analysis

Module scores were computed using the Seurat::AddModuleScore function. This function requires a list of genes comprising the gene expression module to be analyzed. We curated module score gene lists (Supplemental Table S2) from a variety of sources. The ex-vivo activated microglia (exAM) module, which was used to assess artifactual gene expression induced by the single-cell dissociation process, was reported previously (Marsh et al., 2022). We obtained our exAM module list from Table 4 of the Marsh et al. (2022) study. The homeostatic and phagocytic gene module lists were curated from prior scRNA-seq studies of microglia in developing and diseased mouse brain. Two review articles, and the references therein, were particularly useful in curating these lists (Benmamar-Badel et al., 2020; Chen and Colonna, 2021). For the phagocytic list, we only included genes that were strong markers for phagocytic states both during development (Hammond et al., 2019; Li et al., 2019) and in mouse disease models (Benmamar-Badel et al., 2020; Chen and Colonna, 2021). The microglia and macrophage module lists were generated by first comparing microglia and macrophages within our own data, and then refining the lists to ensure they were enriched for published markers of the two cell types in brain (Bennett et al., 2016; Brioschi et al., 2023; Utz et al., 2020; Van Hove et al., 2019).

To identify direct targets of the NRF2 transcription factor, we consulted two reviews and the references therein (Baird and Yamamoto, 2020; Tonelli et al., 2018). We identified 35 genes that met two criteria: 1) There is genetic evidence that they are regulated by NRF2 (i.e. they are upregulated in the context of increased NRF2 activity, and/or downregulated when NRF2 function is impaired; and 2) The genetic locus contains NRF2 binding sites and/or consensus antioxidant-response element (ARE) sequences. This list of 35 genes was used for analysis of Hmox1 cluster marker genes (Fig. 5C). It is important to note that there are far more than 35 genes meeting the first of these criteria; many of these have not been assessed for NRF2 binding or the presence of AREs. Therefore, the spectrum of direct NRF2 targets is probably far larger than the 35 genes we studied here. Our list is simply those that have been validated to date.

### Trajectory analysis

Monocle3 (Cao et al., 2019) was used to perform pseudotime ordering and construct a trajectory describing transcriptional state transitions. To begin, the microglia-only dataset (Fig. 3A) was subsetted to remove minor clusters and the spurious IEG cluster. Remaining clusters after subsetting were: “Homeostatic”, “Apoe”, “Slco2b1”, “Fabp5”, “Apoc1”, “Ccdc152”, “Ccl3/4”, “Spp1”, and “Hmox1.” The subsetted object was then passed through the Seurat dimensionality reduction/clustering pipeline as described above. In the resulting object (Fig. 5D), the cells formerly assigned to the Apoe cluster were now clustered with the Homeostatic and Fabp5 populations. Next, the Seurat object was converted to monocle3 CellDataSet format using the Seurat::as.Cell.Data.Set function. Within monocle3, the cluster_cells function was applied using the existing UMAP reduction as input (clustering resolution was 0.001), followed by the learn_graph function, with learn_graph_control parameter = list(ncenter = 600), which fits a trajectory graph to the data. Finally, to assign a pseudotime value to each cell based on the learned trajectory graph, the monocle3::order_cells function was called. The homeostatic cluster was arbitrarily selected as the root of the trajectory (i.e. pseudotime = 0).

### Immunohistochemistry

Mice were anesthetized and euthanized via decapitation, then eyes were enucleated. Whole eyes were fixed in 4% paraformaldehyde (PFA) in PBS on ice for 1.5 to 2 hours, then washed three times in PBS and stored at 4°C in PBS + 0.02% sodium azide (PBS-azide) until the time of tissue processing. Wholemount retinas were prepared and stained as described previously (Ray et al., 2018). In brief, the anterior eye including lens and vitreous was removed, and the retina was detached out of the eyecup. Retinas were blocked for 2 hours at room temperature in a solution of PBS-azide containing 0.3% Triton X-100 (PBST) and 3% normal donkey serum. Primary antibodies were diluted in blocking solution and incubated for 5-6 days at 4°C with gentle rocking. Retinas were then washed three times in PBS and incubated overnight with secondary antibodies in PBST-azide. Hoechst 33258 was included at this stage as a nuclear counterstain. Additionally, for some experiments, *Griffonia simplicifolia* isolectin B4 (IB4) fluorescent conjugates were included with the primary antibodies to label vasculature. Following secondary antibody staining, retinas were washed at least three times in PBS-azide. To mount stained samples for microscopy, retinas were placed in a dish with PBS and four radial cuts were made, separated by ∼90 degrees. Retinas were mounted ganglion cell-side up on nitrocellulose filters and laid flat with a fine paintbrush. The filter was then placed on a microscope slide and coverslipped with Fluoromount G mounting media (Southern Biotech). The coverslip was sealed with nail polish.

Retinal cryosections were made either from eyecup preparations or from whole eyes. Both procedures have been described previously (Puñal et al., 2019; Ray et al., 2018). In brief, for eyecup sections, the vitreous and lens were removed and the eyecup was cryoprotected in 30% sucrose solution, containing 0.02% sodium azide and PBS, for at least 2 hr. Subsequently the tissue was embedded and frozen in Tissue Freezing Medium (Triangle Biomedical Sciences). For whole eye sections, an elongated cut was made in the cornea to allow for sufficient cryoprotection. Eyes were then sunk in 30% sucrose solution and frozen as described above. Cryosections (20 µm) were collected onto Superfrost Plus slides using a Leica cryostat.

Primary antibodies used in this study are listed in Table 1. All secondary antibodies were acquired from Jackson ImmunoResearch and were conjugated to Alexa-488, Alexa-647, or Cy3 fluorophores.

**Table 1:** Primary antibodies used in this study.

| Antibody | Source | Cat# |
| --- | --- | --- |
| Chicken anti-GFP | ThermoFisher | A10262 |
| Rabbit anti-mCherry (& tdTomato) | Kerafast | EMU106 |
| Rat anti-RFP | ChromoTek | 5f8-20 |
| Rabbit anti-Iba1 | Wako | 019–19741 |
| Goat anti-SPP1/osteopontin | R&D Systems | AF808 |
| Rabbit anti-HMOX1 | Proteintech | 10701-1-AP |
| Mouse anti-ARG1 | Santa Cruz | sc-271430 |
| Rat anti-MHCII-Alexa 488 | BioLegend | 107615 |
| Rat anti-LYVE1 | ThermoFisher | 14-0443-80 |
| Goat anti-CD206 | R&D Systems | AF2535 |
| *Griffonia simplicifolia* IB4 Lectin-Alexa 488 conjugate | ThermoFisher | I21411 |
| *Griffonia simplicifolia* IB4 Lectin-Alexa 647 conjugate | ThermoFisher | I32450 |

### In situ hybridization chain reaction

For in situ hybridization chain reaction (HCR), retinas were dissected in oxygenated AMES media and then fixed overnight in 4% PFA in RNase free PBS. Retinas were washed twice in PBS for 5 min, then dehydrated in methanol and stored at –20°C until the time of tissue processing.

RNase free buffers were prepared for HCR using DEPC-treated H_2_O. Wholemount retinas were rehydrated via a series of MeOH-PBS washes (75:25, 50:50, 25:75, 0:100) before finally incubating in PBST for 1 hour to permeabilize the tissue. To reduce autofluorescence from lipofuscin, tissue was exposed to white LED light (Stillman et al., 2023) as follows: Following rehydration, retinas were placed in a dish with PBS on ice under the LED device (Grow Light CR6000), which was set to 40% brightness for 2 hr. The ice was replaced every 20 min to maintain the temperature close to 4°C.

HCR was performed according to published protocols (Choi et al., 2018) using HCR v3.0 reagents from Molecular Instruments. Gene-specific probes purchased from Molecular Instruments included *Apoc1*, *C1qa*, *Spp1*, *Arg1*, and *Hmox1*. Retinas were permeabilized with 10 µg/mL proteinase K (NEB) for 2 min at room temperature, followed by post-fixation in 4% PFA for 20 min at room temperature and three washes in PBST. Retinas were preconditioned in 30% probe hybridization buffer with rocking on ice, then moved to 37 °C for an additional 30 min of pre-hybridization. Probes were diluted in fresh hybridization buffer at the manufacturer recommended concentration (2 µL per 500 µL hybridization buffer) and retinas were incubated with the probe mixture at 37°C overnight. Subsequently, retinas were washed first with probe wash buffer (Choi et al., 2018), then 5X sodium chloride-sodium citrate buffer + 0.5% Tween (SSCT), then placed in amplification buffer for 30 min at room temperature. Hairpins were prepared for amplification by heating for 90 sec at 95°C, followed by cooling for 30 min at room temperature. Retinas were incubated with hairpins in amplification buffer at 37°C overnight. Retinas were washed in 5X SSCT at room temperature with light protection, then incubated with Hoechst 33258 for 30 min in 5X SSCT and washed again before mounting as described above for immunostained wholemounts. To combine HCR with staining of vasculature, IB4 lectin staining was performed as described in the immunostaining section above after the SSCT wash step, just prior to Hoechst staining and mounting.

### Image acquisition and processing

Nikon AXR and Olympus FV3000 confocal microscopes were used for imaging. Wide fields of view (e.g. Fig. Fig. 2B; Fig. 6A,B,F,G) represent tiled images, acquired either with a 20x air objective or 60x oil objective, and stitched together using Olympus or Nikon software. For cell counting, Z-stacks were acquired from wholemount HCR or immunostained preparations using a 60x oil objective and a Z step size of 1 μm. Laser power and PMT gain were adjusted such that pixels within the brightest cells in a field of view were just below saturation level. This adjustment was especially important to obtain consistent results for HMOX1 antibody staining, because its brightness could vary widely across microglia within an individual retina. For assessment of phagocytic debris, a Z step size of 0.5 µm was used, so as to improve Z resolution of orthogonal XZ or YZ projections

Postprocessing of images was performed in Fiji/ImageJ (Schindelin et al., 2012). All stacks were denoised by applying a median filter with a radius between 0.5 and 2 pixels, applied separately to each channel. Images shown in Figs. 5-8 and associated supplementary figures are Z-projections of the portion of the stack encompassing the RNFL and GCL, or of the cell containing phagocytic debris. For cross-section images (Fig. 2; Fig. 5F), we show Z-projections of the full confocal stack (leaving out any surface slices that are only partially in focus, or which contain acellular material on the section surface). An exception was made in cases where full stack Z-projections would cause two layers of cells present at different stack depths to obscure each other when Z-projected. In these cases, a subset of the stack was Z-projected containing only one of the cell populations. Orthogonal projections of Z stacks were generated using the Fiji Orthogonal Views function. Minor adjustments to brightness and contrast were made in Fiji and/or Adobe Photoshop.

### Quantification and cell counting

To count the fraction of P5 microglia expressing Spp1 and/or Hmox1, the microglia population was sampled by acquiring images from wholemount retinas at standardized locations within central, middle (i.e. mid-peripheral), and peripheral retina, as previously described (O’Sullivan et al., 2017; Perelli et al., 2021; Puñal et al., 2019). In brief, four Z-stacks were acquired at each eccentricity, evenly spaced around concentric circles. Occasionally, it was not possible to acquire all four images at each eccentricity due to tissue damage; however, for each retina used in this analysis, there were at least 3 images at each eccentricity and at least 9 images total. One retina was imaged per animal. Sample sizes (number of animals) are given in figure legends.

To quantify mRNA expression, wholemounts were stained for *C1qa*, *Spp1*, and *Hmox1*. To quantify protein expression, samples were stained for Cx3cr1-GFP; SPP1, and HMOX1. All samples were counterstained with Hoechst nuclear dye, to allow identification of Z slices within the confocal stack that contained the RNFL and GCL. From each Z-stack, we quantified the total number of RNFL/GCL microglia (i.e. *C1qa^+^* or GFP^+^ populations), as well as the fraction of microglia co-expressing the other two markers. RNFL and GCL microglia were combined for this analysis because, due to Z resolution constraints and imperfect retinal flatness, it was difficult to discern whether microglia were contained exclusively within one of these two layers. In any case, we noted that many microglia contacted both layers (see for example Fig. 5H), suggesting that they are a single functional compartment.

Because HMOX1 immunoreactivity varied across a wide dynamic range (Fig. 6B), with the dimmest immunoreactive cells being the most likely to lack *Hmox1* RNA expression (Fig. 6E), we set brightness thresholds for counting cells as HMOX1-positive. To ensure reproducibility in these cell counts, we used consistent image acquisition procedures (see previous section) in which the confocal acquisition settings were optimized for capturing the brightest cells. With this imaging protocol, there was a population of dim cells that could only be discerned with aggressive post-processing brightness/contrast adjustments; such cells were excluded from the HMOX1^+^ cell counts.

For quantification of microglia associated with arteries or veins, we imaged wholemount P5 retinas that had been stained with *Spp1* and *Hmox1* HCR probes together with IB4 lectin to label vasculature. Z-stacks were acquired at the vascular wavefront, in the vicinity of capillaries that were connected closely to a major vessel. Major vessels were identified as arteries based on their thinner appearance and the paucity of capillaries adjacent to them, while veins were recognized by their wider diameter and their close association with nearby capillary loops. The number of *Spp1+* or *Hmox1+* microglia was quantified within each Z-stack as described above.

## ACKNOWLEGEMENTS

This work was supported by the National Eye Institute (EY030611 to J.N.K; EY035119 to J.V-L.; EY5722 to Duke University); by Research to Prevent Blindness (Stein Innovation Award to J.N.K.; unrestricted grant to Duke University); by a Duke University Kahn Neurotechnology award; by a Duke Office of Research & Innovation DST Spark grant; and by the Hartwell Foundation (postdoctoral award to J. V-L). The Duke Flow Cytometry Shared Resource provided essential assistance; and we thank Karen Abramson and the Duke Molecular Genomics Core (Duke Molecular Physiology Institute, Duke University School of Medicine; RRID:SCR_017860) for technical advice and for generating single-cell data. We also thank Ariane Pendragon for mouse colony management; Daniel Saban for helpful discussions; Sarah Hadyniak for the mouse illustration in Fig. 1 and for helpful suggestions; and Caitlin Paisley for numerous helpful comments on the manuscript.

## COMPETING INTERESTS

The authors have no competing interests to disclose.

**Supplemental Table S1**. Top cluster markers: Full dataset

An Excel file showing the top differentially expressed genes in each cluster of the full dataset. Each tab shows the top gene markers enriched in one of the clusters, relative to the rest of the dataset. Genes are sorted by average log2 fold-change. Pct.1, fraction of cells expressing each gene within the cluster of interest; Pct.2, fraction of cells from the rest of the dataset that expresses each gene.

**Supplemental Table S2**. List of genes used for module score analysis

CSV file listing the genes used to define the microglia module; the macrophage (mac) module; the ex vivo activated microglia (ExAM) module; the homeostatic module; and the phagocytic module.

**Supplemental Table S3**. Top cluster markers: Microglia dataset

An Excel file showing the top differentially expressed genes in each cluster of the microglia-only dataset. Layout of tabs and columns as for Supplemental Table S1.

---

**Figure S1.**
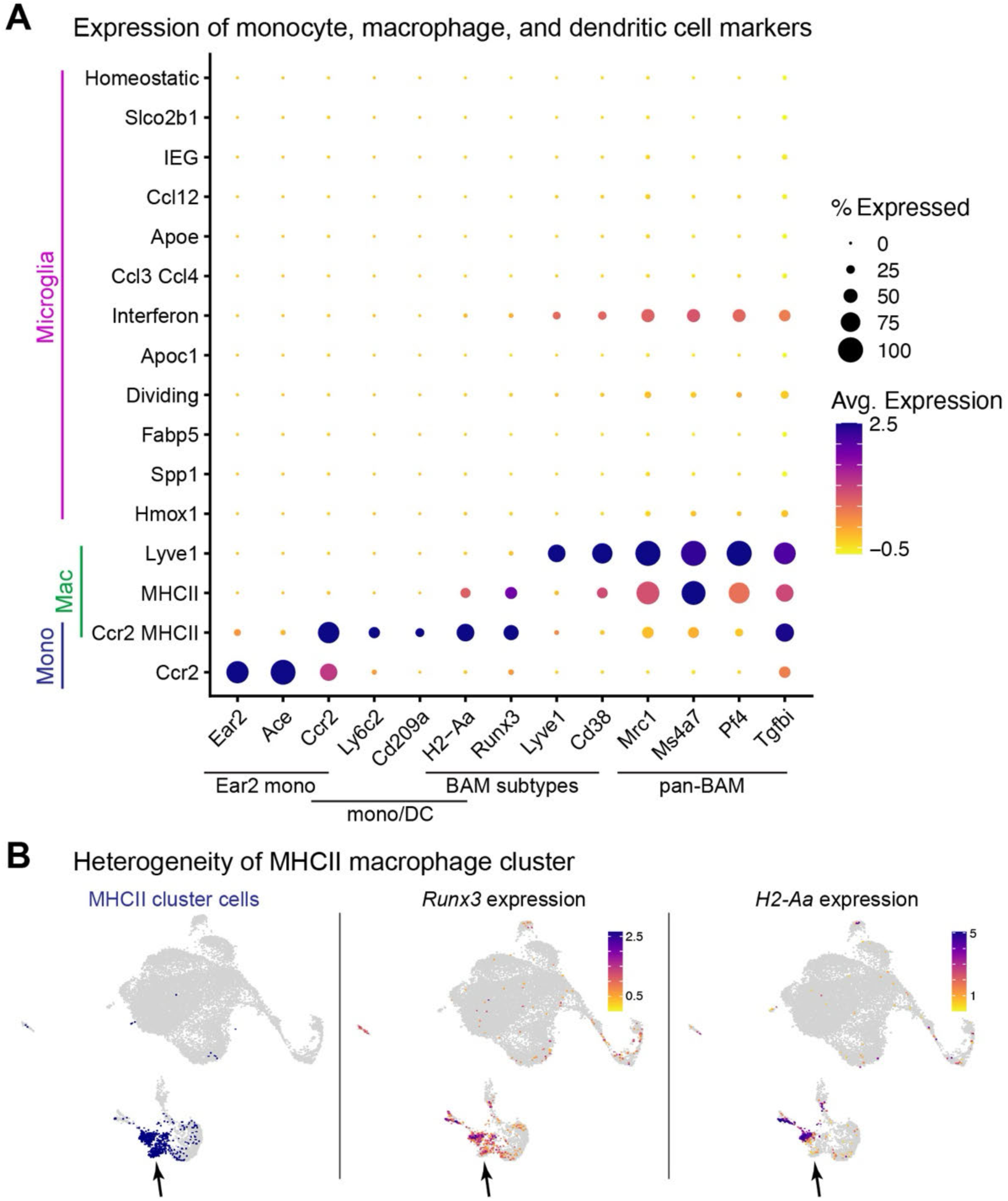
Additional characterization of macrophages and monocytes by scRNA-seq. **A.** Dot plot showing expression of additional monocyte (mono) and macrophage markers, which were used to define the cell types within the non-microglial clusters (see also Fig. 1F and Fig. 3C). Pan-BAM and BAM subset markers from Brioschi et al. (2023); Utz et al. (2020); Van Hove et al. (2019). The Ccr2 cluster expresses markers of *Ear2*+ monocytes (Benhar et al., 2023). The Ccr2-MHCII cluster contains a mixture of different *Ccr2^+^ H2-Aa^+^* cell types. Subsets of this cluster include *Cd209a^+^* dendritic cells (Benhar et al., 2023); *Ly6c2^+^* monocytes or monocyte-derived cells (Van Hove et al., 2019); and MHCII-class BAMs that are most likely immature/differentiating, due to their low expression of pan-BAM markers and their continued *Ccr2* expression. **B.** UMAP plots illustrating heterogeneity of the MHCII macrophage (BAM) cluster. Left, location within UMAP plot of cells assigned to the MHCII cluster (blue). Center and right, Expression of *Runx3* and *H2-Aa*. Two distinct groups of MHCII cluster cells are evident: a *Runx3^+^ H2-Aa^−^* population (arrow), and a *Runx3^+^ H2-Aa^+^*double-positive population.

**Figure S2:**
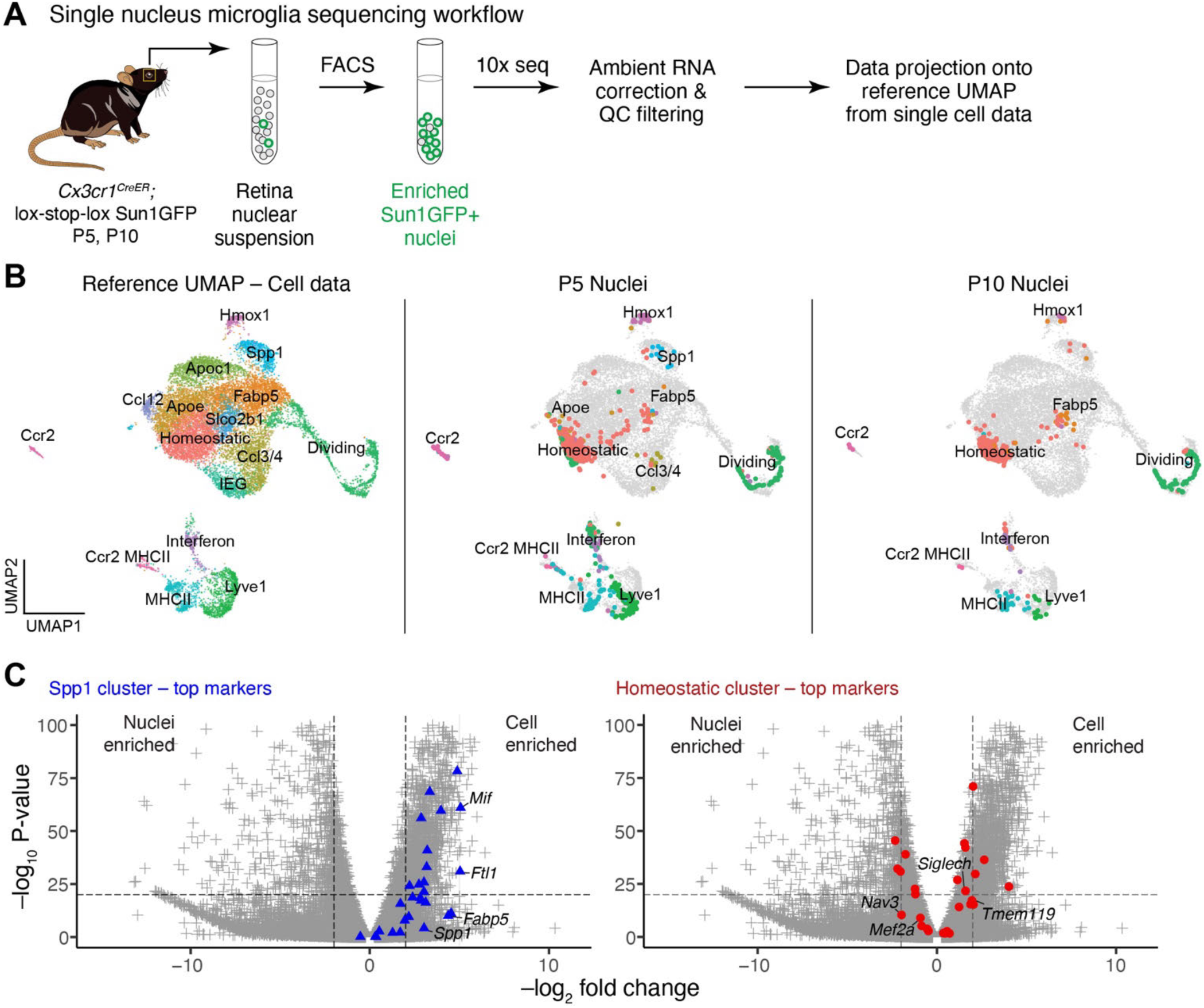
Single nucleus RNA-seq of retinal myeloid cells. **A.** Workflow for isolation and single-nucleus (sn) RNA-seq of GFP-labeled myeloid cell nuclei at P5 and P10. **B.** P5 and P10 snRNA-seq data projected onto the reference UMAP generated from single-cell data. Left panel reproduces the reference UMAP from Fig. 1C, with cluster labels. Reference UMAP is gray in center, right panels. Colored dots in center, right panels show projected single nuclei with colors matching the cluster assignment as in left panel. Clusters such as Homeostatic, Hmox1, Ccr2, and the macrophage clusters were detected in both cell and nucleus datasets. However, many other scRNA-seq clusters were underrepresented or absent in snRNA-seq data, including IEG and Spp1. **C.** Pseudobulk analysis comparing gene expression between the single-cell and single-nucleus datasets. Volcano plots show that markers of the Spp1 cluster (blue, top) were systematically underrepresented in the nucleus data, explaining poor detection of this microglial state via snRNA-seq. By contrast, no method-dependent bias was observed for markers of the homeostatic cluster (red, bottom).

**Figure S3.**
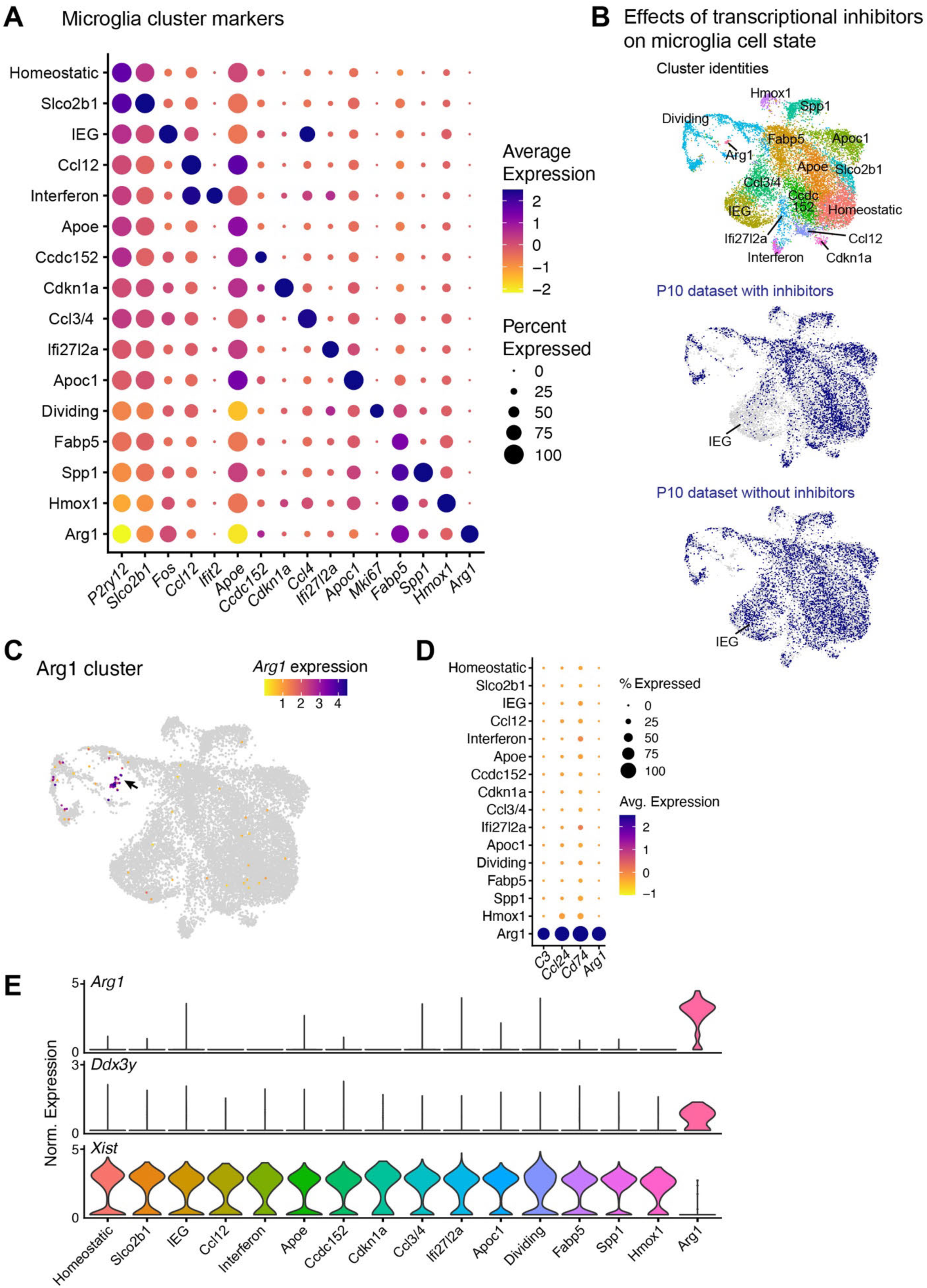
Additional scRNA-seq characterization of microglia clusters. **A.** Dot plot generated from microglia-only dataset (B, left) showing expression of a selected top marker of each microglial cluster. **B.** Comparison of UMAP locations for cells from the P10 dataset generated without transcriptional inhibitors (center) and the P10 dataset generated with inhibitors (bottom). UMAP plot showing microglia cluster identities (from Fig. 3A) is shown at top for reference. Cells from the inhibitor dataset are largely absent from the IEG cluster (labeled). However, inhibitor-treated cells populate all other regions of the UMAP. The Ccl3/4 cluster was diminished in the inhibitor sample, but *Ccl3*^+^ and *Ccl4*^+^ microglia have previously been shown to exist in retina and brain (Marsh et al., 2022; Anderson et al., 2022). Neither P10 dataset populated the Arg1 cluster, which was specific to P5 microglia (Fig. 3C). **E.** Arg1 cluster (arrow) is the smallest of the 16 microglia clusters (n = 35 cells). *Arg1* expression is highly selective for this cluster. **F.** Dot plot showing expression of top Arg1 cluster markers across microglia dataset. **G.** Violin plots showing expression of sex-specific transcripts across clusters. Arg1 cluster is enriched for Y chromosome transcripts (*Ddx3y*) and depleted for the female-specific transcript *Xist*, suggesting that most microglia within this cluster derive from males.

**Figure S4.**
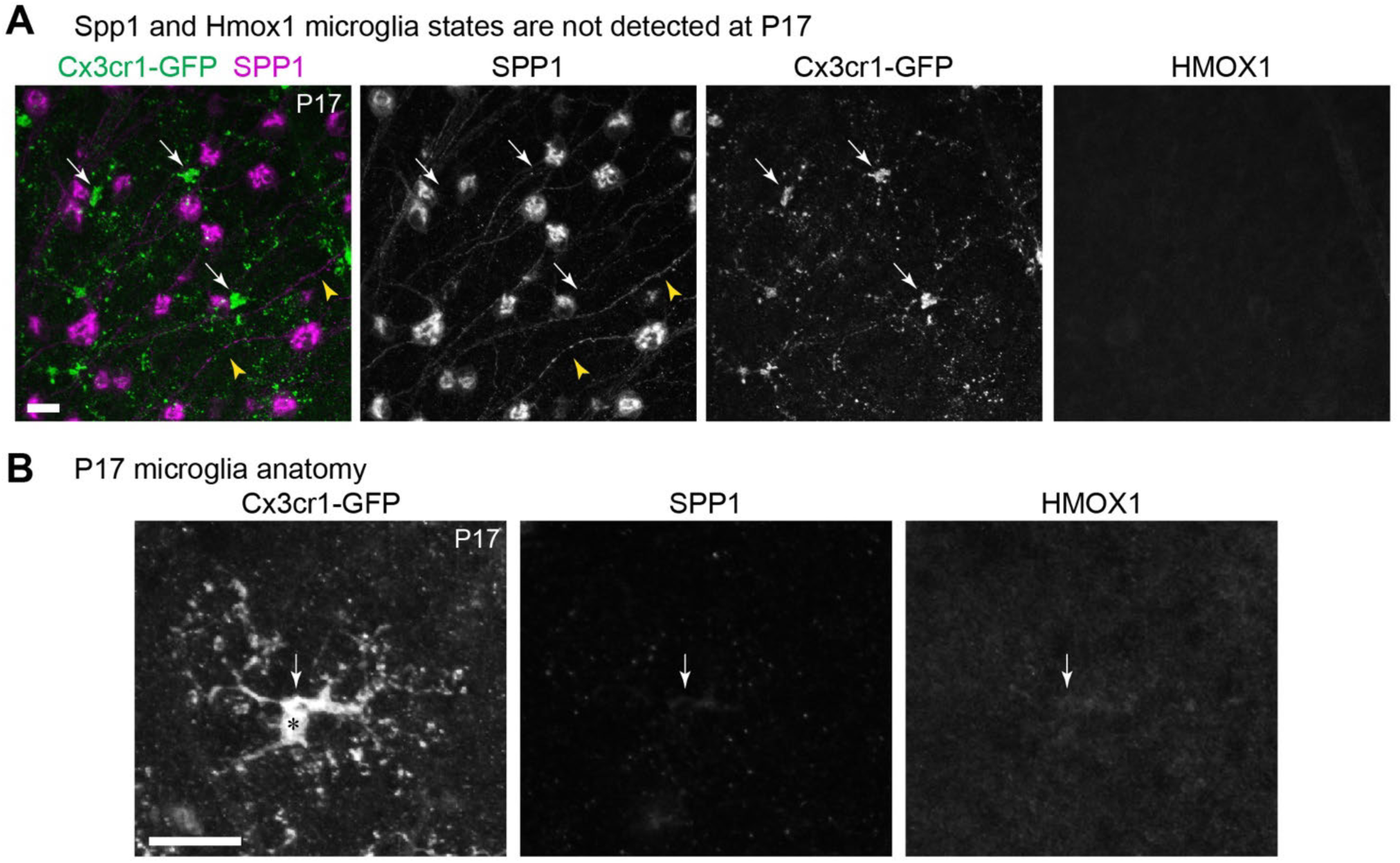
Additional phenotyping of retinal microglia at P5 and P17. **A.** Triple immunolabeling at P17 for microglia (Cx3cr1-GFP), SPP1, and HMOX1. As illustrated by this image (representative of n = 4 retinas from 4 mice), HMOX1 and SPP1 were not detected in microglia (arrows). SPP1 is instead expressed by RGCs at P17 (arrowheads indicate SPP^+^ RGC axons). **B.** Representative example of P17 microglial cell anatomy. Microglia at this age exhibit characteristics of homeostatic microglia, including a highly ramified morphology and absence of SPP1 and HMOX1 expression.

## REFERENCES

Anderson, S.R., J.M. Roberts, N. Ghena, E.A. Irvin, J. Schwakopf, I.B. Cooperstein, A. Bosco, and M.L. Vetter. 2022. Neuronal apoptosis drives remodeling states of microglia and shifts in survival pathway dependence. Elife. 11.

Anderson, S.R., J.M. Roberts, J. Zhang, M.R. Steele, C.O. Romero, A. Bosco, and M.L. Vetter. 2019. Developmental Apoptosis Promotes a Disease-Related Gene Signature and Independence from CSF1R Signaling in Retinal Microglia. Cell Rep. 27:2002–2013.e2005.

Anderson, S.R., and M.L. Vetter. 2019. Developmental roles of microglia: A window into mechanisms of disease. Dev. Dyn. 248:98–117.

Baird, L., and M. Yamamoto. 2020. The Molecular Mechanisms Regulating the KEAP1-NRF2 Pathway. Mol. Cell. Biol. 40.

Barclay, K.M., N. Abduljawad, Z. Cheng, M.W. Kim, L. Zhou, J. Yang, J. Rustenhoven, J.A. Mazzitelli, L.C.D. Smyth, D. Kapadia, S. Brioschi, W. Beatty, J. Hou, N. Saligrama, M. Colonna, G. Yu, J. Kipnis, and Q. Li. 2024. An inducible genetic tool to track and manipulate specific microglial states reveals their plasticity and roles in remyelination. Immunity. 57:1394–1412 e1398.

Benhar, I., J. Ding, W. Yan, I.E. Whitney, A. Jacobi, M. Sud, G. Burgin, K. Shekhar, N.M. Tran, C. Wang, Z. He, J.R. Sanes, and A. Regev. 2023. Temporal single-cell atlas of non-neuronal retinal cells reveals dynamic, coordinated multicellular responses to central nervous system injury. Nat. Immunol. 24:700–713.

Benmamar-Badel, A., T. Owens, and A. Wlodarczyk. 2020. Protective Microglial Subset in Development, Aging, and Disease: Lessons From Transcriptomic Studies. Front Immunol. 11:430.

Bennett, M.L., F.C. Bennett, S.A. Liddelow, B. Ajami, J.L. Zamanian, N.B. Fernhoff, S.B. Mulinyawe, C.J. Bohlen, A. Adil, A. Tucker, I.L. Weissman, E.F. Chang, G. Li, G.A. Grant, M.G. Hayden Gephart, and B.A. Barres. 2016. New tools for studying microglia in the mouse and human CNS. Proceedings of the National Academy of Sciences. 113:E1738–E1746.

Bisht, K., K.A. Okojie, K. Sharma, D.H. Lentferink, Y.Y. Sun, H.R. Chen, J.O. Uweru, S. Amancherla, Z. Calcuttawala, A.B. Campos-Salazar, B. Corliss, L. Jabbour, J. Benderoth, B. Friestad, W.A. Mills, 3rd, B.E. Isakson, M.E. Tremblay, C.Y. Kuan, and U.B. Eyo. 2021. Capillary-associated microglia regulate vascular structure and function through PANX1-P2RY12 coupling in mice. Nat Commun. 12:5289.

Brioschi, S., J.A. Belk, V. Peng, M. Molgora, P.F. Rodrigues, K.M. Nguyen, S. Wang, S. Du, W.L. Wang, G.E. Grajales-Reyes, J.M. Ponce, C.M. Yuede, Q. Li, J.M. Baer, D.G. DeNardo, S. Gilfillan, M. Cella, A.T. Satpathy, and M. Colonna. 2023. A Cre-deleter specific for embryo-derived brain macrophages reveals distinct features of microglia and border macrophages. Immunity.

Cao, J., M. Spielmann, X. Qiu, X. Huang, D.M. Ibrahim, A.J. Hill, F. Zhang, S. Mundlos, L. Christiansen, F.J. Steemers, C. Trapnell, and J. Shendure. 2019. The single-cell transcriptional landscape of mammalian organogenesis. Nature. 566:496–502.

Chen, Y., and M. Colonna. 2021. Microglia in Alzheimer’s disease at single-cell level. Are there common patterns in humans and mice? J. Exp. Med. 218.

Choi, H.M.T., M. Schwarzkopf, M.E. Fornace, A. Acharya, G. Artavanis, J. Stegmaier, A. Cunha, and N.A. Pierce. 2018. Third-generation in situ hybridization chain reaction: multiplexed, quantitative, sensitive, versatile, robust. Development. 145.

Escoubas, C.C., L.C. Dorman, P.T. Nguyen, C. Lagares-Linares, H. Nakajo, S.R. Anderson, J.J. Barron, S.D. Wade, B. Cuevas, I.D. Vainchtein, N.J. Silva, R. Guajardo, Y. Xiao, P.V. Lidsky, E.Y. Wang, B.M. Rivera, S.E. Taloma, D.K. Kim, E. Kaminskaya, H. Nakao-Inoue, B. Schwer, T.D. Arnold, A.B. Molofsky, C. Condello, R. Andino, T.J. Nowakowski, and A.V. Molofsky. 2024. Type-I-interferon-responsive microglia shape cortical development and behavior. Cell. 187:1936–1954 e1924.

Fourgeaud, L., P.G. Través, Y. Tufail, H. Leal-Bailey, E.D. Lew, P.G. Burrola, P. Callaway, A. Zagórska, C.V. Rothlin, A. Nimmerjahn, and G. Lemke. 2016. TAM receptors regulate multiple features of microglial physiology. Nature. 532:240–244.

Ghena, N., S.R. Anderson, J.M. Roberts, E. Irvin, J. Schwakopf, A. Bosco, and M.L. Vetter. 2025. CD11c-Expressing Microglia Are Transient, Driven by Interactions With Apoptotic Cells. Glia.

Giera, S., R. Luo, Y. Ying, S.D. Ackerman, S.-J. Jeong, H.M. Stoveken, C.J. Folts, C.A. Welsh, G.G. Tall, B. Stevens, K.R. Monk, and X. Piao. 2018. Microglial transglutaminase-2 drives myelination and myelin repair via GPR56/ADGRG1 in oligodendrocyte precursor cells. eLife. 7.

Gray, D.A., and J. Woulfe. 2005. Lipofuscin and aging: a matter of toxic waste. Sci Aging Knowledge Environ. 2005:re1.

Hagemeyer, N., K.M. Hanft, M.A. Akriditou, N. Unger, E.S. Park, E.R. Stanley, O. Staszewski, L. Dimou, and M. Prinz. 2017. Microglia contribute to normal myelinogenesis and to oligodendrocyte progenitor maintenance during adulthood. Acta Neuropathol. 134:441–458.

Hammond, T.R., C. Dufort, L. Dissing-Olesen, S. Giera, A. Young, A. Wysoker, A.J. Walker, F. Gergits, M. Segel, J. Nemesh, S.E. Marsh, A. Saunders, E. Macosko, F. Ginhoux, J. Chen, R.J.M. Franklin, X. Piao, S.A. McCarroll, and B. Stevens. 2019. Single-Cell RNA Sequencing of Microglia throughout the Mouse Lifespan and in the Injured Brain Reveals Complex Cell-State Changes. Immunity. 50:253–271.e256.

Hammond, T.R., D. Robinton, and B. Stevens. 2018. Microglia and the Brain: Complementary Partners in Development and Disease. Annu. Rev. Cell. Dev. Biol. 34:523–544.

He, D., H. Xu, H. Zhang, R. Tang, Y. Lan, R. Xing, S. Li, E. Christian, Y. Hou, P. Lorello, B. Caldarone, J. Ding, L. Nguyen, D. Dionne, P. Thakore, A. Schnell, J.R. Huh, O. Rozenblatt-Rosen, A. Regev, and V.K. Kuchroo. 2021. Disruption of the IL-33-ST2-AKT signaling axis impairs neurodevelopment by inhibiting microglial metabolic adaptation and phagocytic function. Immunity.

Ishii, T., K. Itoh, S. Takahashi, H. Sato, T. Yanagawa, Y. Katoh, S. Bannai, and M. Yamamoto. 2000. Transcription factor Nrf2 coordinately regulates a group of oxidative stress-inducible genes in macrophages. J. Biol. Chem. 275:16023–16029.

Jiang, D., C.A. Burger, V. Akhanov, J.H. Liang, R.D. Mackin, N.E. Albrecht, P. Andrade, D.P. Schafer, and M.A. Samuel. 2022. Neuronal signal-regulatory protein alpha drives microglial phagocytosis by limiting microglial interaction with CD47 in the retina. Immunity. 55:2318–2335 e2317.

Krasemann, S., C. Madore, R. Cialic, C. Baufeld, N. Calcagno, R. El Fatimy, L. Beckers, E. O’Loughlin, Y. Xu, Z. Fanek, D.J. Greco, S.T. Smith, G. Tweet, Z. Humulock, T. Zrzavy, P. Conde-Sanroman, M. Gacias, Z. Weng, H. Chen, E. Tjon, F. Mazaheri, K. Hartmann, A. Madi, J.D. Ulrich, M. Glatzel, A. Worthmann, J. Heeren, B. Budnik, C. Lemere, T. Ikezu, F.L. Heppner, V. Litvak, D.M. Holtzman, H. Lassmann, H.L. Weiner, J. Ochando, C. Haass, and O. Butovsky. 2017. The TREM2-APOE Pathway Drives the Transcriptional Phenotype of Dysfunctional Microglia in Neurodegenerative Diseases. Immunity. 47:566–581 e569.

Kubota, Y., K. Takubo, T. Shimizu, H. Ohno, K. Kishi, M. Shibuya, H. Saya, and T. Suda. 2009. M-CSF inhibition selectively targets pathological angiogenesis and lymphangiogenesis. The Journal of experimental medicine. 206:1089–1102.

Lawrence, A.R., A. Canzi, C. Bridlance, N. Olivie, C. Lansonneur, C. Catale, L. Pizzamiglio, B. Kloeckner, A. Silvin, D.A.D. Munro, A. Fortoul, D. Boido, F. Zehani, H. Cartonnet, S. Viguier, G. Oller, P. Squarzoni, A. Candat, J. Helft, C. Allet, F. Watrin, J.B. Manent, P. Paoletti, D. Thieffry, L. Cantini, C. Pridans, J. Priller, A. Gelot, P. Giacobini, L. Ciobanu, F. Ginhoux, M.S. Thion, L. Lokmane, and S. Garel. 2024. Microglia maintain structural integrity during fetal brain morphogenesis. Cell. 187:962–980 e919.

Li, Q., Z. Cheng, L. Zhou, S. Darmanis, N.F. Neff, J. Okamoto, G. Gulati, M.L. Bennett, L.O. Sun, L.E. Clarke, J. Marschallinger, G. Yu, S.R. Quake, T. Wyss-Coray, and B.A. Barres. 2019. Developmental Heterogeneity of Microglia and Brain Myeloid Cells Revealed by Deep Single-Cell RNA Sequencing. Neuron. 101:207–223.e210.

Marsh, S.E. 2021. Custom Visualizations & Functions for Streamlined Analyses of Single Cell Sequencing.

Marsh, S.E., A.J. Walker, T. Kamath, L. Dissing-Olesen, T.R. Hammond, T.Y. de Soysa, A.M.H. Young, S. Murphy, A. Abdulraouf, N. Nadaf, C. Dufort, A.C. Walker, L.E. Lucca, V. Kozareva, C. Vanderburg, S. Hong, H. Bulstrode, P.J. Hutchinson, D.J. Gaffney, D.A. Hafler, R.J.M. Franklin, E.Z. Macosko, and B. Stevens. 2022. Dissection of artifactual and confounding glial signatures by single-cell sequencing of mouse and human brain. Nat. Neurosci. 25:306–316.

Masuda, T., R. Sankowski, O. Staszewski, C. Böttcher, L. Amann, Sagar, C. Scheiwe, S. Nessler, P. Kunz, G. van Loo, V.A. Coenen, P.C. Reinacher, A. Michel, U. Sure, R. Gold, D. Grün, J. Priller, C. Stadelmann, and M. Prinz. 2019. Spatial and temporal heterogeneity of mouse and human microglia at single-cell resolution. Nature. 566:388–392.

Mendiola, A.S., J.K. Ryu, S. Bardehle, A. Meyer-Franke, K.K. Ang, C. Wilson, K.M. Baeten, K. Hanspers, M. Merlini, S. Thomas, M.A. Petersen, A. Williams, R. Thomas, V.A. Rafalski, R. Meza-Acevedo, R. Tognatta, Z. Yan, S.J. Pfaff, M.R. Machado, C. Bedard, P.E. Rios Coronado, X. Jiang, J. Wang, M.A. Pleiss, A.J. Green, S.S. Zamvil, A.R. Pico, B.G. Bruneau, M.R. Arkin, and K. Akassoglou. 2020. Transcriptional profiling and therapeutic targeting of oxidative stress in neuroinflammation. Nat. Immunol. 21:513–524.

Mendiola, A.S., Z. Yan, K. Dixit, J.R. Johnson, M. Bouhaddou, A. Meyer-Franke, M.G. Shin, Y. Yong, A. Agrawal, E. MacDonald, G. Muthukumar, C. Pearce, N. Arun, B. Cabriga, R. Meza-Acevedo, M. Alzamora, S.S. Zamvil, A.R. Pico, J.K. Ryu, N.J. Krogan, and K. Akassoglou. 2023. Defining blood-induced microglia functions in neurodegeneration through multiomic profiling. Nat. Immunol. 24:1173–1187.

Mo, A., E.A. Mukamel, F.P. Davis, C. Luo, G.L. Henry, S. Picard, M.A. Urich, J.R. Nery, T.J. Sejnowski, R. Lister, S.R. Eddy, J.R. Ecker, and J. Nathans. 2015. Epigenomic Signatures of Neuronal Diversity in the Mammalian Brain. Neuron. 86:1369–1384.

Mondo, E., S.C. Becker, A.G. Kautzman, M. Schifferer, C.E. Baer, J. Chen, E.J. Huang, M. Simons, and D.P. Schafer. 2020. A Developmental Analysis of Juxtavascular Microglia Dynamics and Interactions with the Vasculature. J Neurosci. 40:6503–6521.

Moreno-Garcia, A., A. Kun, O. Calero, M. Medina, and M. Calero. 2018. An Overview of the Role of Lipofuscin in Age-Related Neurodegeneration. Front Neurosci. 12:464.

Munro, D.A.D., K. Movahedi, and J. Priller. 2022. Macrophage compartmentalization in the brain and cerebrospinal fluid system. Sci Immunol. 7:eabk0391.

Nemes-Baran, A.D., D.R. White, and T.M. DeSilva. 2020. Fractalkine-Dependent Microglial Pruning of Viable Oligodendrocyte Progenitor Cells Regulates Myelination. Cell Rep. 32:108047.

O’Koren, E.G., C. Yu, M. Klingeborn, A.Y.W. Wong, C.L. Prigge, R. Mathew, J. Kalnitsky, R.A. Msallam, A. Silvin, J.N. Kay, C. Bowes Rickman, V.Y. Arshavsky, F. Ginhoux, M. Merad, and D.R. Saban. 2019. Microglial Function Is Distinct in Different Anatomical Locations during Retinal Homeostasis and Degeneration. Immunity. 50:723–737.e727.

O’Sullivan, M.L., V.M. Puñal, P.C. Kerstein, J.A. Brzezinski, T. Glaser, K.M. Wright, and J.N. Kay. 2017. Astrocytes follow ganglion cell axons to establish an angiogenic template during retinal development. Glia. 65:1697–1716.

Paisley, C.E., and J.N. Kay. 2021. Seeing stars: Development and function of retinal astrocytes. Dev. Biol. 478:144–154.

Parkhurst, C.N., G. Yang, I. Ninan, J.N. Savas, J.R. Yates, J.J. Lafaille, B.L. Hempstead, D.R. Littman, and W.-B. Gan. 2013. Microglia Promote Learning-Dependent Synapse Formation through Brain-Derived Neurotrophic Factor. Cell. 155:1596–1609.

Perelli, R.M., M.L. O’Sullivan, S. Zarnick, and J.N. Kay. 2021. Environmental oxygen regulates astrocyte proliferation to guide angiogenesis during retinal development. Development. 148:dev199418.

Puñal, V.M., C.E. Paisley, F.S. Brecha, M.A. Lee, R.M. Perelli, J. Wang, E.G. O’Koren, C.R. Ackley, D.R. Saban, B.E. Reese, and J.N. Kay. 2019. Large-scale death of retinal astrocytes during normal development is non-apoptotic and implemented by microglia. PLoS Biol. 17:e3000492.

Rajesh, A., S. Droho, and J.A. Lavine. 2022. Macrophages in close proximity to the vitreoretinal interface are potential biomarkers of inflammation during retinal vascular disease. J Neuroinflammation. 19:203.

Ransohoff, R.M., and V.H. Perry. 2009. Microglial physiology: unique stimuli, specialized responses. Annu. Rev. Immunol. 27:119–145.

Ray, T.A., S. Roy, C. Kozlowski, J. Wang, J. Cafaro, S.W. Hulbert, C.V.E. Wright, G.D. Field, and J.N. Kay. 2018. Formation of retinal direction-selective circuitry initiated by starburst amacrine cell homotypic contact. eLife. 7:e34241.

Rosmus, D.D., J. Koch, A. Hausmann, A. Chiot, F. Arnhold, T. Masuda, K. Kierdorf, S.M. Hansen, H. Kuhrt, J. Froba, J. Wolf, S. Boneva, M. Gericke, B. Ajami, M. Prinz, C. Lange, and P. Wieghofer. 2024. Redefining the ontogeny of hyalocytes as yolk sac-derived tissue-resident macrophages of the vitreous body. J Neuroinflammation. 21:168.

Saederup, N., A.E. Cardona, K. Croft, M. Mizutani, A.C. Cotleur, C.L. Tsou, R.M. Ransohoff, and I.F. Charo. 2010. Selective chemokine receptor usage by central nervous system myeloid cells in CCR2-red fluorescent protein knock-in mice. PLoS One. 5:e13693.

Schindelin, J., I. Arganda-Carreras, E. Frise, V. Kaynig, M. Longair, T. Pietzsch, S. Preibisch, C. Rueden, S. Saalfeld, B. Schmid, J.-Y. Tinevez, D.J. White, V. Hartenstein, K. Eliceiri, P. Tomancak, and A. Cardona. 2012. Fiji: an open-source platform for biological-image analysis. Nat. Methods. 9:676–682.

Somebang, K., J. Rudolph, I. Imhof, L. Li, E.C. Niemi, J. Shigenaga, H. Tran, T.M. Gill, I. Lo, B.A. Zabel, G. Schmajuk, B.T. Wipke, S. Gyoneva, L. Jandreski, M. Craft, G. Benedetto, E.D. Plowey, I. Charo, J. Campbell, C.J. Ye, S.S. Panter, M.C. Nakamura, W. Eckalbar, and C.L. Hsieh. 2021. CCR2 deficiency alters activation of microglia subsets in traumatic brain injury. Cell Rep. 36:109727.

Stillman, J.M., F. Mendes Lopes, J.P. Lin, K. Hu, D.S. Reich, and D.P. Schafer. 2023. Lipofuscin-like autofluorescence within microglia and its impact on studying microglial engulfment. Nat Commun. 14:7060.

Stratoulias, V., R. Ruiz, S. Kanatani, A.M. Osman, L. Keane, J.A. Armengol, A. Rodriguez-Moreno, A.N. Murgoci, I. Garcia-Dominguez, I. Alonso-Bellido, F. Gonzalez Ibanez, K. Picard, G. Vazquez-Cabrera, M. Posada-Perez, N. Vernoux, D. Tejera, K. Grabert, M. Cheray, P. Gonzalez-Rodriguez, E.M. Perez-Villegas, I. Martinez-Gallego, A. Lastra-Romero, D. Brodin, J. Avila-Carino, Y. Cao, M. Airavaara, P. Uhlen, M.T. Heneka, M.E. Tremblay, K. Blomgren, J.L. Venero, and B. Joseph. 2023. ARG1-expressing microglia show a distinct molecular signature and modulate postnatal development and function of the mouse brain. Nat. Neurosci. 26:1008–1020.

Stuart, T., A. Butler, P. Hoffman, C. Hafemeister, E. Papalexi, W.M. Mauck, Y. Hao, M. Stoeckius, P. Smibert, and R. Satija. 2019. Comprehensive Integration of Single-Cell Data. Cell. 177:1888–1902.e1821.

Tammela, T., G. Zarkada, H. Nurmi, L. Jakobsson, K. Heinolainen, D. Tvorogov, W. Zheng, C.a. Franco, A. Murtomäki, E. Aranda, N. Miura, S. Ylä-Herttuala, M. Fruttiger, T. Mäkinen, A. Eichmann, J.W. Pollard, H. Gerhardt, and K. Alitalo. 2011. VEGFR-3 controls tip to stalk conversion at vessel fusion sites by reinforcing Notch signalling. Nat. Cell Biol. 13:1202–1213.

Thrupp, N., C. Sala Frigerio, L. Wolfs, N.G. Skene, N. Fattorelli, S. Poovathingal, Y. Fourne, P.M. Matthews, T. Theys, R. Mancuso, B. de Strooper, and M. Fiers. 2020. Single-Nucleus RNA-Seq Is Not Suitable for Detection of Microglial Activation Genes in Humans. Cell Rep. 32:108189.

Tirosh, I., B. Izar, S.M. Prakadan, M.H. Wadsworth, 2nd, D. Treacy, J.J. Trombetta, A. Rotem, C. Rodman, C. Lian, G. Murphy, M. Fallahi-Sichani, K. Dutton-Regester, J.R. Lin, O. Cohen, P. Shah, D. Lu, A.S. Genshaft, T.K. Hughes, C.G. Ziegler, S.W. Kazer, A. Gaillard, K.E. Kolb, A.C. Villani, C.M. Johannessen, A.Y. Andreev, E.M. Van Allen, M. Bertagnolli, P.K. Sorger, R.J. Sullivan, K.T. Flaherty, D.T. Frederick, J. Jane-Valbuena, C.H. Yoon, O. Rozenblatt-Rosen, A.K. Shalek, A. Regev, and L.A. Garraway. 2016. Dissecting the multicellular ecosystem of metastatic melanoma by single-cell RNA-seq. Science. 352:189–196.

Tonelli, C., I.I.C. Chio, and D.A. Tuveson. 2018. Transcriptional Regulation by Nrf2. Antioxid. Redox Signal. 29:1727–1745.

Trapnell, C., D. Cacchiarelli, J. Grimsby, P. Pokharel, S. Li, M. Morse, N.J. Lennon, K.J. Livak, T.S. Mikkelsen, and J.L. Rinn. 2014. The dynamics and regulators of cell fate decisions are revealed by pseudotemporal ordering of single cells. Nat. Biotechnol. 32:381–386.

Unlu, G., B. Prizer, R. Erdal, H.W. Yeh, E.C. Bayraktar, and K. Birsoy. 2022. Metabolic-scale gene activation screens identify SLCO2B1 as a heme transporter that enhances cellular iron availability. Mol. Cell. 82:2832–2843 e2837.

Utz, S.G., P. See, W. Mildenberger, M.S. Thion, A. Silvin, M. Lutz, F. Ingelfinger, N.A. Rayan, I. Lelios, A. Buttgereit, K. Asano, S. Prabhakar, S. Garel, B. Becher, F. Ginhoux, and M. Greter. 2020. Early Fate Defines Microglia and Non-parenchymal Brain Macrophage Development. Cell. 181:557–573.e518.

Van Hove, H., L. Martens, I. Scheyltjens, K. De Vlaminck, A.R. Pombo Antunes, S. De Prijck, N. Vandamme, S. De Schepper, G. Van Isterdael, C.L. Scott, J. Aerts, G. Berx, G.E. Boeckxstaens, R.E. Vandenbroucke, L. Vereecke, D. Moechars, M. Guilliams, J.A. Van Ginderachter, Y. Saeys, and K. Movahedi. 2019. A single-cell atlas of mouse brain macrophages reveals unique transcriptional identities shaped by ontogeny and tissue environment. Nat. Neurosci. 22:1021–1035.

Witcher, K.G., C.E. Bray, T. Chunchai, F. Zhao, S.M. O’Neil, A.J. Gordillo, W.A. Campbell, D.B. McKim, X. Liu, J.E. Dziabis, N. Quan, D.S. Eiferman, A.J. Fischer, O.N. Kokiko-Cochran, C. Askwith, and J.P. Godbout. 2021. Traumatic Brain Injury Causes Chronic Cortical Inflammation and Neuronal Dysfunction Mediated by Microglia. J Neurosci. 41:1597–1616.

Wlodarczyk, A., I.R. Holtman, M. Krueger, N. Yogev, J. Bruttger, R. Khorooshi, A. Benmamar-Badel, J.J. Boer-Bergsma, N.A. Martin, K. Karram, I. Kramer, E.W. Boddeke, A. Waisman, B.J. Eggen, and T. Owens. 2017. A novel microglial subset plays a key role in myelinogenesis in developing brain. The EMBO Journal. 36:3292–3308.

Young, M.D., and S. Behjati. 2020. SoupX removes ambient RNA contamination from droplet-based single-cell RNA sequencing data. Gigascience. 9.

